# Adhesion-Controlled Mechanics of the Glial Niche Regulate Neural Stem Cell Proliferative Potential

**DOI:** 10.1101/2024.12.11.628019

**Authors:** Anna Segú Cristina, David Briand, Aman Kukde, Emeline Perthame, Stéphane Rigaud, Jean-Yves Tinevez, Agata Banach-Latapy, Nick D.L. Owens, Léo Valon, Yohanns Bellaïche, Pauline Spéder

## Abstract

Controlled proliferation of neural stem cells (NSCs) builds a functional nervous system during development. While their cellular niche is recognized as a signalling hub, the contribution of its structure and mechanics in regulating neurogenesis remains unexplored. The *Drosophila* larval central nervous system contains self-renewing NSCs in close contact with cortex glial cells. Transcriptomics identified a triad of immunoglobulin superfamily cell adhesion molecules (Dpr10/Dpr6 in glia and DIP-α in NSCs) which physically and mechanically connect the NSC and glial membranes, acting as mechanoregulators. Their disruption increases glial cortical tension, causing non-autonomous mitotic defects in NSCs, characterized by abnormal spindle morphologies and impaired mitotic progression. Additionally, elevated glial tensile forces increase Lamin content in NSCs, a protective response also resulting in nuclear deformation. Ultimately NSC proliferative potential and genome integrity are compromised. Our study reveals that the native mechanical properties of the niche are transmitted to NSCs and regulate their function.

**Graphical abstract:** 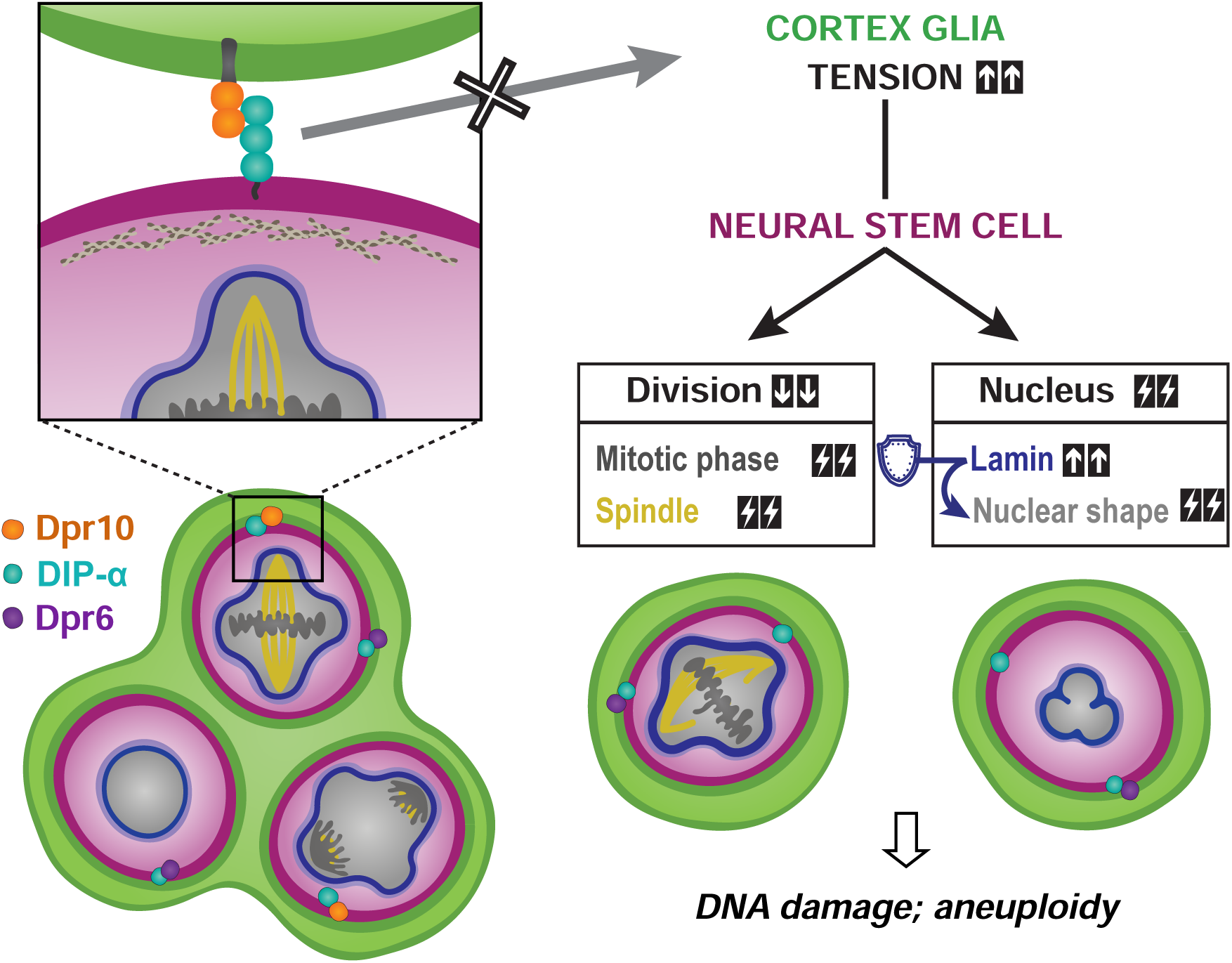

## INTRODUCTION

Development of a central nervous system (CNS) relies on the precise generation of diverse neurons, which later form functional networks. This task is devolved to neural stem cells (NSCs), multipotent progenitors that self-renew while producing differentiated progeny^1,2^. Tight regulation of their proliferation ensures the timely production of the right number and types of neurons. Such control is primarily fueled by intrinsic regulators, including asymmetric division machinery and fate determinants^1^. NSCs also integrate external cues, ranging from local signals to systemic information^3,4^. A key provider and modulator of these external influences is the specific cellular microenvironment where NSCs reside, the niche^5,6^. The neurogenic niche is heterogeneous and includes the NSCs themselves, glial cells, resident immune cells, blood vessels and extracellular matrix components. In adults, the niche is known for providing paracrine and soluble signals that regulate NSC behaviour^7–9^. Beyond signalling, the niche is a structural scaffold rich in cell interactions organizing a complex architecture. For example, NSCs directly contact blood vessels, which are also covered by a meshwork of glial end feet^5,8^. Diverse cell-cell recognition and adhesion molecules support these interactions^10,11^, like integrins linking to the extracellular matrix and N-cadherin forming adherens junctions between NSCs. Removing these molecules alters NSC behaviour and neurogenesis, yet it is unclear whether it is a result of their role as signal transducer, of their architectural functions, or of both^10–13^. Although the architecture of the neurogenic niche is a striking feature, its influence on neurogenesis, whether at the level of individual NSCs or population-wide, has been scarcely explored. Similarly, the nature and role of mechanical forces generated and exerted by niche cells, tied to adhesive properties, are poorly understood, despite the CNS undergoing morphogenetic changes both during development and ageing, implying a crucial role for tissue mechanics^14–16^. Ultimately, how structural and mechanical inputs from cellular interactions within the native niche directly regulate NSC proliferation *in vivo* remains unclear.

The *Drosophila* CNS has become a valuable *in vivo* model for studying neurogenesis, thanks to its conservation of key cellular mechanisms, compelling genetics, and ease of manipulation^17,18^. *Drosophila* NSCs originate from the neuroectoderm during embryonic development, dividing to produce primary neurons^2,17^. After a period of quiescence at the end of embryogenesis, NSCs reactivate in a nutrient-dependent manner and proliferate during larval stages to generate secondary neurons, which form most of the adult CNS. These larval NSCs exist in a multilayered cellular niche, comprising a glial subtype called the cortex glia (CG). CG signalling is essential for neurogenesis at multiple levels. They regulate NSC division in physiological conditions^19–23^, protect NSCs from external biochemical stresses^24,25^ and control the survival, positioning and branching of newborn neurons^20,26,27^. CG form a striking architecture around NSCs, encasing each NSC lineage in a bespoke membrane chamber while spanning the CNS as a network ^26,28,29^ (Figure 1A). This structure develops alongside NSC proliferative behaviour: CG cells first expand and send membranes, driven by nutritional input, then enclose NSCs at the time of their mitotic re-entry, and finally keep adapting as NSC lineages grow^23,26,27,30,31^ (Figure 1B). Thus, proliferating NSCs are enclosed in close-fitting CG membrane chambers, which form a grid-type pattern at the level of the NSC population.

**Figure 1.**
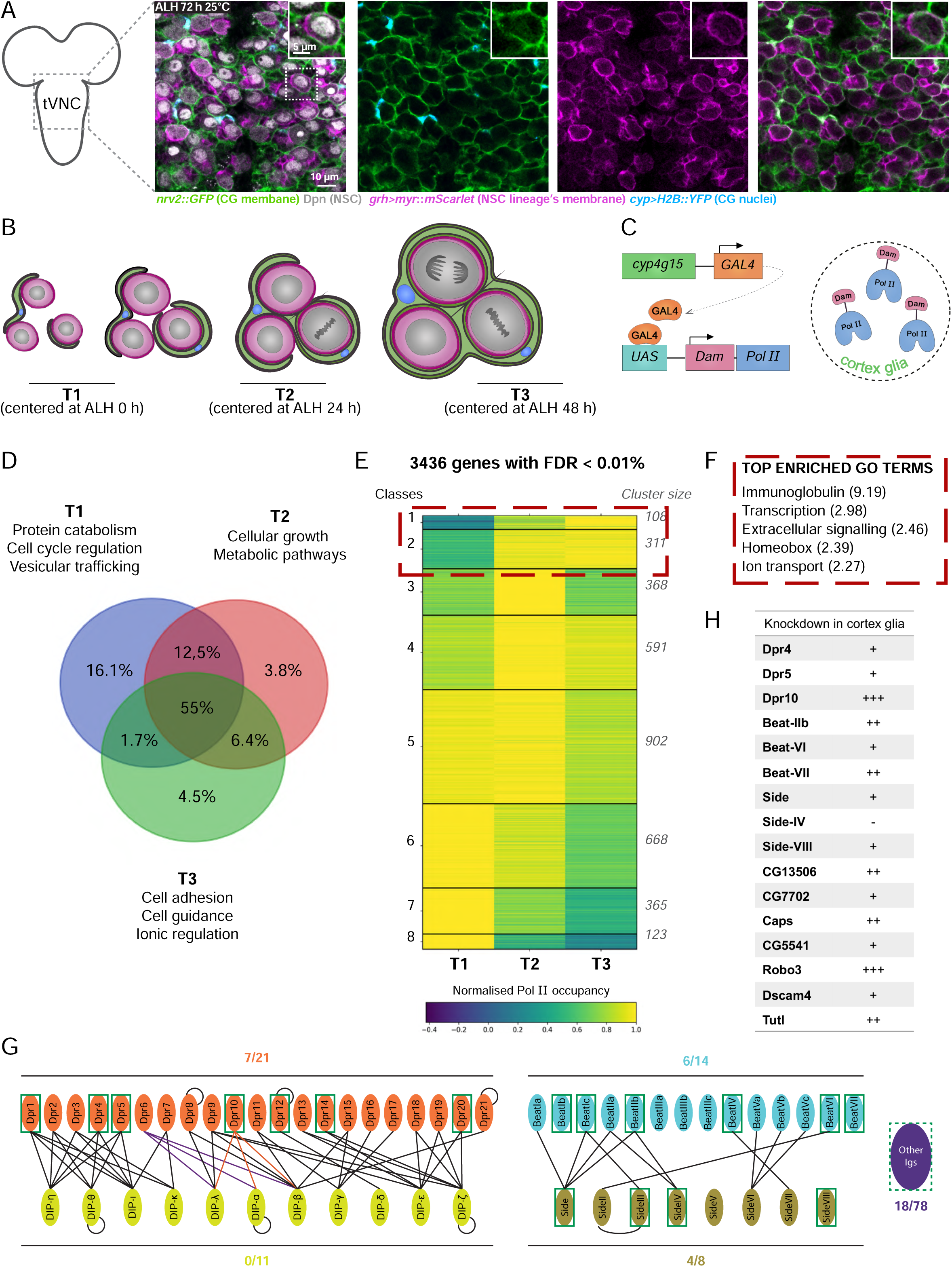
Transcriptional analysis of the cortex glia overtime identifies the immunoglobulin superfamily as a key player in in niche morphogenesis. (A) Close up of a thoracic ventral nerve cord (tVNC) from a *Drosophila* larva showing CG tightly encasing NSC lineages, forming a grid of membrane chambers across the NSC population. CG membranes are marked with *nrv2::GFP* and nuclei with *cyp4g15-GAL4* (*cyp*) driving *H2B∷YFP*. NSC membranes are labelled with *grh-GAL4* driving *myr-mScarlet* and nuclei with Dpn antibody. (B) Schematic of CG morphogenesis through larval stages, showing three time windows: (T1) pre-CG membrane growth; (T2) during CG membrane growth; (T3) during NSC encasement by CG membranes. ALH: after larval hatching. (C) CG specific profiling of RNA Pol II occupancy in the genome (TaDa-PollI) can be obtained by expressing Dam-Pol II under the control of the *cyp4g15-GAL4* driver in a time-controlled manner thanks to the UAS/GAL4 system coupled with a thermosensitive version of the GAL80 repressor. (D) GO term analysis of unique genes in each time window revealed enriched categories with relevant biological functions for niche formation. (E-F) 3436 genes were identified (FDR < 0.01%) from the TaDa-PolII and grouped into 8 clusters based on expression profiles. Clusters 1 and 2 with increasing expression overtime revealed enrichment in members of the immunoglobulin superfamily (IgSF). Normalised PolII occupancy is scaled by the maximum occupancy of each gene, such that the maximum value for each gene is set to 1. (G) Extracellular interactome of the *Drosophila* IgSF. Main subfamilies include Dpr (21 members) interacting with DIP (11 members), and Beat (14 members) with Side (8 members). Left numbers and boxes indicate the number and identity of members found in the TaDa-PolII. (H) Summary table of the functional screening of genes with increasing expression over time. *cyp4g15-GAL4* was crossed with specific RNAi lines and CG grid-type structure and NSC numbers and proliferation were assessed qualitatively. The number of "+" indicates the strengths of CG and NSC phenotypes, irrespective of nature. See also Table S2 and Figure S1.

How are niche and NSC physical interactions controlled during neurodevelopment? Leveraging this model of NSC niche, we show that the mechanical properties of the cellular niche regulate NSC function. A triad of cell adhesion molecules of the immunoglobulin superfamily (Dpr10/Dpr6 in the CG and DIP-α in NSCs) bridges the NSC and CG membranes and control CG tensile forces. Disrupting these complexes increases CG cortical tension, leading to mitotic defects in NSCs, including abnormal spindle morphologies and impaired mitotic progression. NSCs also exhibit variable levels of DNA damage and aneuploidy. In parallel, increased tension in the CG drives Lamin accumulation in NSC nuclei, causing nuclear deformation yet protective of mitosis. These findings highlight the importance of the NSC microenvironment’s mechanical properties in regulating NSC function.

## RESULTS

### Transcriptional analysis of cortex glia during niche morphogenesis identifies a strong enrichment in the immunoglobulin superfamily during niche formation

To investigate the interactions between NSC and CG during chamber formation, we assessed gene expression in CG along this process (Figure 1B). We used the Targeted DamID Pol II technique^32,33^, which allows the profiling of gene expression in a specific cell type and at chosen timepoints without the need for cell isolation (Figures 1C and S1A). This method uses a DNA adenine methyltransferase (Dam) fused to a core RNA polymerase II (Pol II) subunit to methylate GATC sequences at Pol II binding sites, indicating paused and active transcription. Controlling its expression by the GAL4/GAL80/UAS system enables time- and cell-specific methylation signals, which are mapped to identify genes enriched for Pol II binding^34^. DamID Pol II was expressed under the control of a CG specific GAL4 driver (*cyp4g15-GAL4*^26,27^), and restricted to three time windows covering the main steps of chamber formation: T1, before nutrition-induced membrane growth; T2, during membrane growth; and T3, at the time of NSC encasing (Figure S1B). Annotation of the genes specific to T1, T2 and T3 (Figure 1D) revealed gene enrichments, with functions pertaining to cellular growth at T2, and guidance and adhesion pathways at T3. Further clustering of these data identified genes with different temporal expression patterns (Figure 1E and Table S2). Genes with increasing expression over time (classes 1 and 2) revealed a striking enrichment in members of the immunoglobulin superfamily (IgSF, Figure 1F). We found that out of 35 IgSF members expressed in CG (Figure 1G), 26 were increasing overtime.

IgSF members are cell-cell adhesion molecules (CAMs) which form homophilic or heterophilic complexes with other members of the superfamily^35,36^. These IgSF-CAMs harbour the most frequent extracellular domain in humans and participate in many biological processes requiring cell interaction and communication, including migration, adhesion, guidance and self-recognition^37^. In the mammalian CNS, they are important mediators of neuronal connections and neurogenesis^38,39^. The *Drosophila* IgSF comprises 130 non-antibody, non-T cell receptor members and is also crucial in the assembly of neural circuitry, by mediating the targeting and anchoring of opposing synaptic partners through the generation of unique combinations of cell interactions^40–43^. Fly IgSF-CAMs are divided in four main subfamilies determined by sequence similarity and interactome^44,45^ (Figure 1G). The DIP subfamily interacts with the Dpr subfamily, and the Beat subfamily with the Side subfamily. Interestingly, while CG express several Dprs, they do not express any member of the DIP subfamily (Figure 1G), suggesting that Dpr and DIP must be mediating interactions between CG and other cell types.

We performed a functional screen on these 26 IgSF-CAM candidates, knocking them down specifically in the CG. We scored qualitatively for defects in CG membrane structure (formation of individual chambers around NSC and of the grid-type pattern) using the CG marker *nrv2::GFP* (Figure 1A) and in NSC number and proliferation with combined staining of the NSC-specific transcription factor Deadpan (Dpn, Figure 1A) with the mitotic marker phospho-Histone 3 (pH3). Scoring was done at mid-L3 larval stage (60-72 h After Larval Hatching (ALH)), when the CG structure is formed and NSCs actively divide, focusing solely on the thoracic Ventral Nerve Cord (tVNC) for simplicity. This region and timeframe will be consistently used throughout this study. Our qualitative screen uncovered a variety of phenotypes across all tested candidates, revealing an essential and multifaceted role of IgSF-CAMs in niche morphogenesis (Figure 1H). Although most defects affected CG structure, *dpr10* knockdown in CG produced a distinct phenotype in NSCs.

### Loss of *dpr10* in the cortex glia alters the nuclear shape, proliferation and mitotic progression of neural stem cells in a cell non-autonomous manner

While *dpr10* knockdown in the CG did not lead to detectable changes in CG grid-type structure, it caused a striking alteration of nuclear shapes in NSCs, overtly visible in z-projections (Figure 2A, dashed yellow outlines). We relied on Dpn as a proxy for nuclear shape during interphase/early prophase, as it remains within the nuclear envelope during these stages, only becoming cytoplasmic from prometaphase through cytokinesis (Figure S2A). This readout revealed that, instead of the rounder, oblong typical form of control interphase and prophase nuclei, NSC nuclei under *dpr10* knockdown in CG displayed a range of contorted shapes (Figures 2A-B), at a penetrance of 20 % (Figure 2C). Of note, we investigated the use of several fully automated approaches based on well-established geometric parameters (*e.g.*, roundness, sphericity, convexity) to get finer details on nuclei morphology. However, these geometric parameters were insufficient to reliably and automatically detect changes for small and varied 3D indentations, and phenotypic strength will be assessed qualitatively.

**Figure 2.**
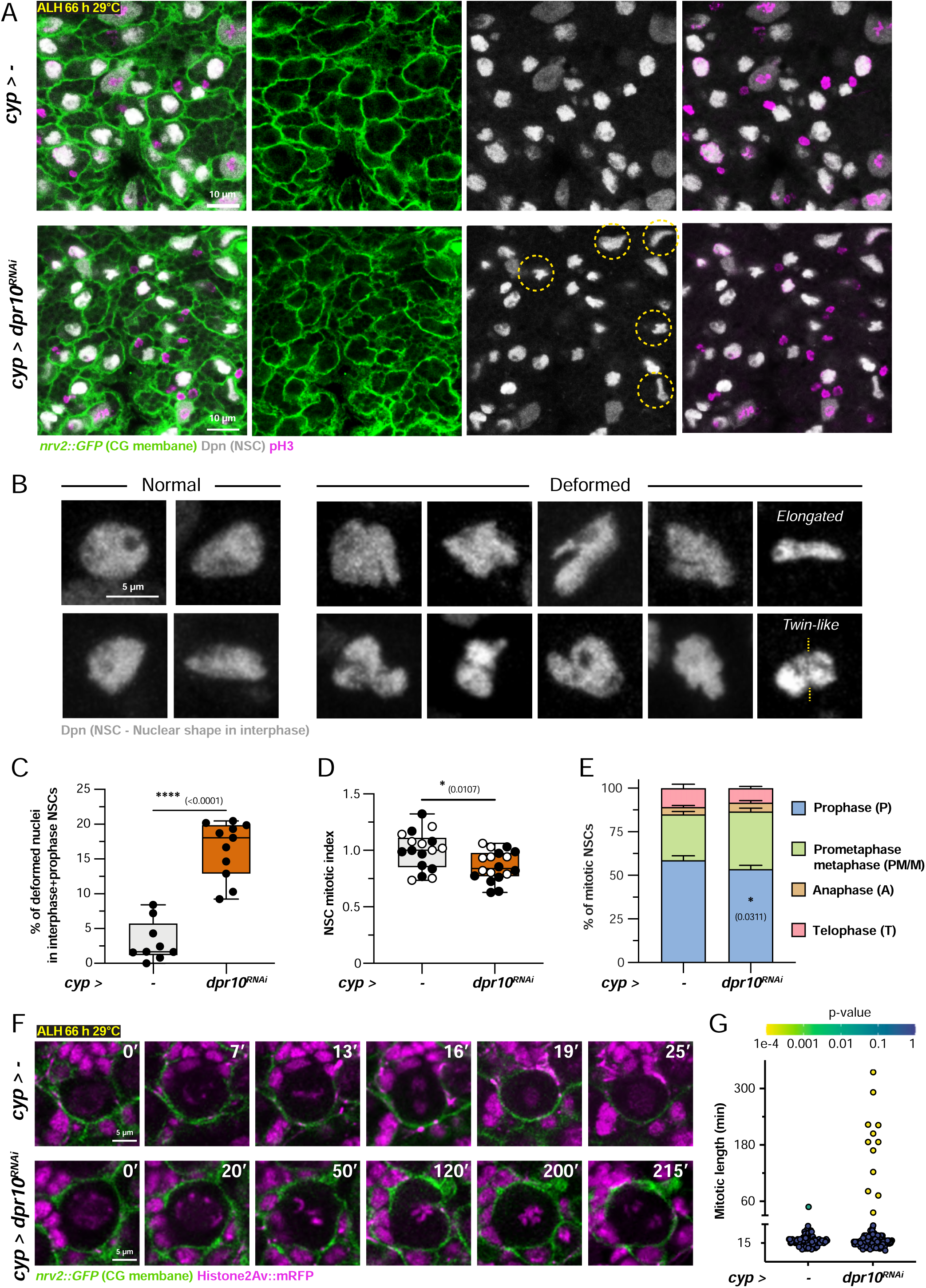
*dpr10* is required non-autonomously in the cortex glia to control neural stem cell proliferation and nuclear shape. (A) Confocal images of the tVNC showing CG membrane and NSC division in control condition (*cyp > -*) and under *dpr10* knockdown in the CG (*cyp > dpr10 RNAi^VDRC103511^*). Dashed yellow circles indicate examples of deformed nuclei. (B) Close up of NSC nuclei illustrating the range of normal and deformed shapes in interphase/early prophase. Dpn stays within the nuclear lamina in these phases, during which it serves as a proxy for nuclear shape. The yellow dash line marks the axis from which the nucleus’ halves look mirror image from each other. (C) Box plot of the percentage of NSCs (in interphase + early prophase) showing deformed nuclear shapes in *cyp > -* (n = 9 tVNCs) and *cyp > dpr10 RNAi^VDRC103511^* (n = 11 tVNCs). Unpaired Student t-test. (D-E) Box plot of mitotic index (normalized to control) and stacked bar chart of phase distribution in NSCs for *cyp > -* (n = 17 tVNCs) and *cyp > dpr10 RNAi^VDRC103511^* (n = 18 tVNCs). Unpaired Student t-test (D) and two-way ANOVA with Šídák multiple comparison test (E). (F) Still images of a time-lapse movie of mitotic NSCs in control (*cyp > -*, Movie S1) and *dpr10* knockdown in the CG (*cyp > dpr10 RNAi^VDRC103511^*, Movie S2). Some NSCs from *cyp > dpr10 RNAi^VDRC103511^* were arrested in metaphase for over more than 1h. (G) Graph of the probability of each measured mitotic length to be significantly different from the control distribution. *cyp > -* (n = 98 NSCs) and *cyp > dpr10 RNAi^VDRC103511^* (n = 162 NSCs). 6.8% of NSCs in *cyp > dpr10 RNAi^VDRC103511^* showed a significant disruption of the cell cycle progression. For box plots, individual values are superimposed and colour-coded to represent multiple experimental replicates. ns, p≥0.05; *, p < 0.05; **, p <0.01; ***, p<0.001; ****, p<0.0001 P: Prophase, PM/M: Prometaphase/metaphase, A: Anaphase, T: Telophase. See also Table S2 and Figure S2.

We assayed NSC division at the same mid-L3 stage, during which NSCs undergo mitosis almost every hour at their peak rate^46,47^. Using pH3 staining, we primarily found that *dpr10* knockdowns in CG resulted in a decrease of NSC mitotic index compared to control (Figure 2D), as well as in a mild, yet significant, change in the proportion of the different mitotic steps, with a shift towards prometaphase/metaphase (Figure 2E), suggesting that NSCs might spend longer in these phases. NSC numbers per tVNC were not changed (Figure S2B). Interestingly, accumulated biological replicates of *dpr10* knockdown in CG revealed variability in phenotypic strength, with some exhibiting a higher mitotic index and more pronounced shifts in mitotic phase distribution (*e.g.*, increased prophase or prometaphase/metaphase) (Figure S2C-D). The elevated mitotic index likely results from prolonged duration in certain phases, making the proportions of pH3^+^ cells higher. These shifts support the hypothesis that NSCs may linger or even arrest in these phases, potentially slowing the mitotic process. Then, ultimately NSCs might fail to enter division either because of repetitive damages or of stochastic stronger *dpr10* downregulation.

To further characterize NSCs’ mitotic phenotype under *dpr10* knockdown in the CG, we used live imaging to record NSC divisions with fluorescently tagged Histone2Av (*His2Av::mRFP*). While all NSCs analyzed in control were able to complete mitosis in an average of 17 min, several NSCs experienced very long metaphases, ranging from 1 h up to 5 h under *dpr10* knockdown in the CG (Figures 2F-G and Movies S1-S3). Altogether, these data show that knockdown of *dpr10* in the CG alters NSC mitotic progression.

Since NSCs divide in an asymmetric fashion, we assessed the localization of the basal determinant Miranda, required for proper asymmetric segregation of fate determinants during mitotis^48,49^ and did not observe differences under *dpr10* knockdown in CG compared to control (Figure S2E). Similarly, the non-muscular molecular motor Myosin II, essential both for asymmetric polarization and cytokinesis^50,51^, displayed the well-documented bottleneck pattern during telophase and the formation of a midbody (Figure S2F), visualized with a GFP-tagged Myosin II regulatory light chain under its native promoter (*sqh::GFP*)^52^. The lack of supernumerary NSCs, revealing of symmetric division^53,54^, is consistent with these findings (Figure S2B). Thus, asymmetric machinery does not seem affected in NSCs under *dpr10* knockdown in the CG.

Taken together, our results show that *dpr10* functions in the CG to regulate NSC nuclear shape, mitotic progression and proliferation.

### Dpr10 is required in the cortex glia for membrane contact with the neural stem cells

How can Dpr10 in the CG regulate NSC proliferation in a non-autonomous manner? IgSF-CAMs support cell-to-cell interactions through their binding properties to specific partners. We postulated that Dpr10 could influence NSC function by providing adhesion between NSCs and CG, bringing their membranes in close contact with each other. The GFP Reconstitution Across Synaptic Partners (GRASP) system^55^ uses two complementary fragments of GFP (Split-GFP^56^) on transmembrane proteins. Fluorescence is restored when the fragments are close, indicating membrane proximity and likely cell contact. We used parallel genetic binary systems (GAL4/UAS and LexO/LexAOp) to express one GFP fragment in the CG and the complementary other in NSCs (Figure 3A). In control condition, we were able to detect a continuous and mostly uniform GFP signal overlapping with a reporter for NSC membrane (Figure 3B). Conversely, the GFP signal under *dpr10* knockdown in CG appeared discontinuous and heterogenous, with spots of brighter signal, compared to NSC membrane marker. We quantified NSC-CG proximity by measuring the ratios between the GFP signal and the NSC membrane in both conditions and found that it was in overall decreased when *dpr10* was knocked down in the CG (Figure 3C). In addition, we noticed that the morphology of the NSC membrane appeared altered, with more angular contours and inward indentations. Geometric measurements of NSC membrane contour at the equatorial plane (respective to Dpn staining) revealed an increased convexity (Figure 3D). These results show that Dpr10-mediated interactions within the niche are required for proper contact between NSC and CG membranes and for NSC cell shape. They further suggest that Dpr10 binds to a partner on the NSC membrane.

**Figure 3.**
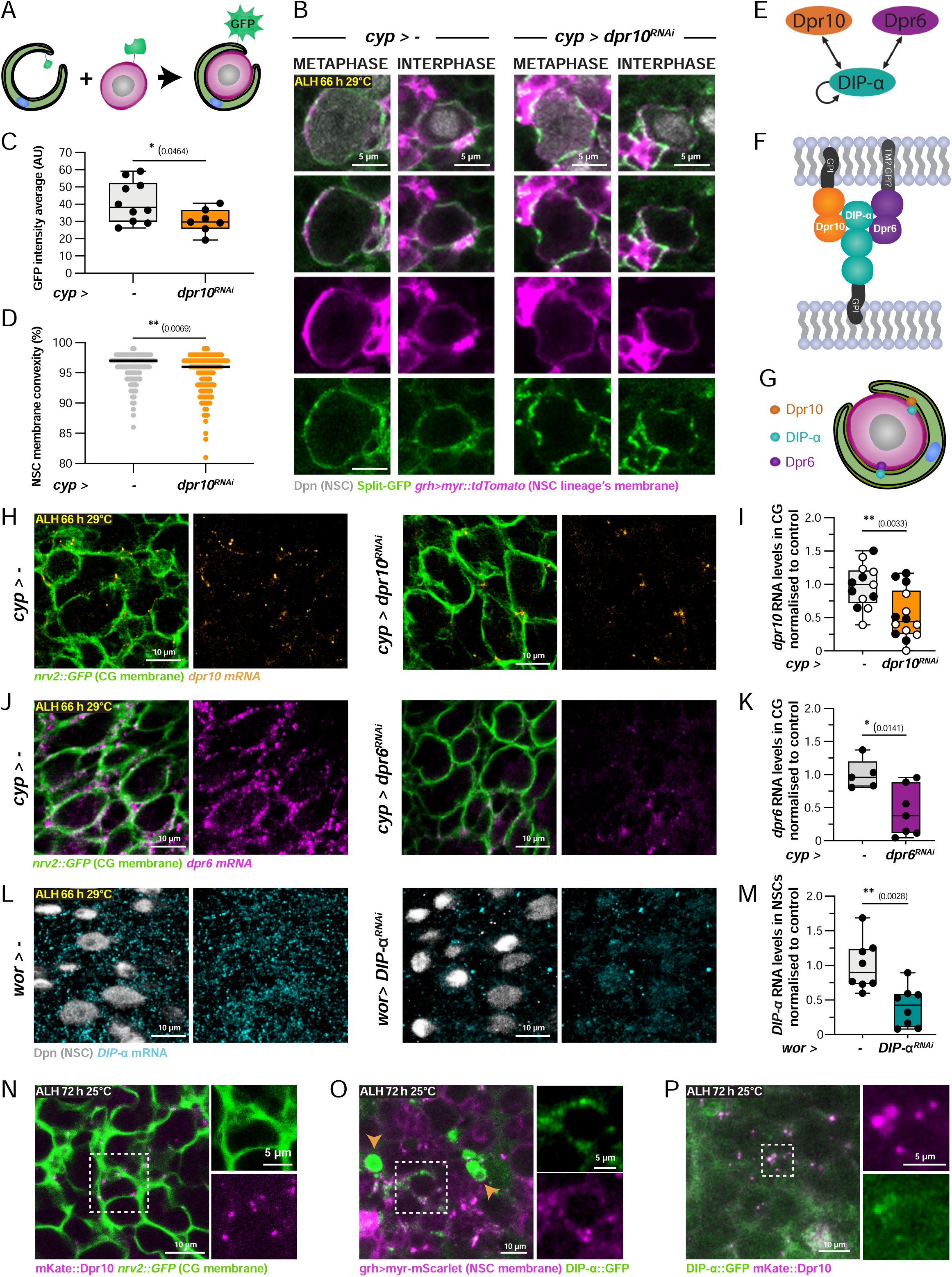
Dpr10 maintains cortex glia and neural stem cell membranes’ proximity and is expressed in the neurogenic niche as an IgSF-CAM triad with DIP-α and Dpr6. (A) Sketch of the split-GFP technique. One of the two GFP fragments is expressed in CG and the complementary one is expressed in NSCs. The contact and assembly of the two fragments reconstitutes GFP fluorescence, making it a readout of membrane proximity and cell contact. (B) Confocal images of NSCs expressing split-GFP (green) between CG and NSCs. *cyp4g15-GAL4* drove *Dpr10* RNAi and one GFP fragment in CG. *grh-LexA* drove the complementary fragment of the GFP and a membrane marker (*myr-tomato)* in NSCs. (C) Box plot of split-GFP intensity ratio in NSC membranes in *cyp >-* (n=10 tVNCs) and *cyp > dpr10 RNAi^VDRC103511^* (n = 7 tVNCs). Unpaired Student’s t-test. (D) Dot plot of membrane convexity in NSCs in *cyp > -* (n = 11 tVNCs and 197 NSCs) and *cyp > dpr10 RNAi^VDRC103511^*(n = 7 tVNCs and 126 NSCs). Dot = NSC. Unpaired Student t-test on average values per tVNC. (E) Schematics of the IgSF-CAM triad. Dpr10 binds to DIP-α, which can also bind to Dpr6 and exhibits homophilic interactions with other DIP-α proteins. (F) DIP-α and Dpr10 anchor to the membrane using a GPI group, while Dpr6 might use a transmembrane domain. They interact through their Ig domains. (G) Schematics of the potential cellular interactions in the niche through the IgSF-CAM triad. (H-I) Confocal images and box plot of *dpr10* RNA (orange) levels in the CG for *cyp>-* (n = 13 tVNCs) and *cyp> dpr10 RNAi^VDRC103511^*(n = 14 tVNCs). Unpaired Student’s t-test. (J-K) Confocal images and box plot of *dpr6* RNA (magenta) levels in the CG for *cyp>-* (n = 5 tVNCs) and *cyp> dpr6 RNAi^VDRC41161^*(n = 7 tVNCs). Unpaired Student’s t-test. (L-M) Confocal images and box plot of *DIP-α* RNA (cyan) levels in NSCs for *wor>-* (n = 8 tVNCs) and *wor UAS-DIP-α RNAi^VDRC104044^* (n = 8 tVNCs). Unpaired Student’s t-test. (N) Still image of a time-lapse movie (Movie S4) of a CRISPR/Cas9 engineered line *mKate2::dpr10* (magenta). CG membrane: *Nrv2::GFP* (green). Insert highlights the expression of mKate2::dpr10 over the CG membrane. (O) Still image of a time-lapse movie (Movie S6) of a CRISPR/Cas9 engineered line *DIP-α::GFP* (green). NSC and progeny membrane: *grh^D4^-myr-mScarlet* (magenta). *DIP-α::GFP* is expressed in neurons (orange arrowheads) and colocalizes with the NSC membrane. Insert highlights the expression of DIP-α::GFP over the NSC membrane. (P) Still image of a time-lapse movie (Movie S7) of the CRISPR/Cas9 *mKate2::dpr10* (magenta) and *DIP-α::GFP* (green) together. Insert highlights the partial co-localization between these two fusion proteins. For box plots, individual values are superimposed and colour-coded to represent multiple experimental replicates. ns, p≥0.05; *, p < 0.05; **, p <0.01; ***, p<0.001; ****, p<0.0001 See also Table S2 and Figure S3.

### DIP-α, a binding partner of Dpr10 and of Dpr6, is expressed in neural stem cells, while Dpr10 and Dpr6 are both expressed in the cortex glia

The “Dpr-ome” interaction network currently comprises 21 Dpr proteins interacting in a complex pattern with 11 DIPs^40–42,44,57^. Most Dprs bind to multiple DIPs and conversely. Previous biochemical and biological approaches^40,43,44^ have shown that Dpr10 establishes heterophilic interactions with DIP-α. DIP-α can also interact with Dpr6 as well as set up homophilic interactions with itself (Figures 3E-F). DIP-α has three Ig domains like all DIPs, while Dpr10 and Dpr6, like all Dprs, have two. DIPs and Dprs are GPI-anchored^57^, likely requiring transmembrane coreceptors, and thus predicted to be cell surface glycoproteins^58^. GPI-anchoring has been confirmed for Dpr10 and DIP-α, while Dpr6 may have a transmembrane domain (Figure 3F).

A hypothesis would see DIP-α as the partner of Dpr10, and possibly Dpr6, in NSCs (Figure 3G). The DIP-α/Dpr10/Dpr6 triad is implicated in *Drosophila* CNS development, particularly through heterophilic interactions crucial for neuronal targeting and synaptic pairing in the visual system^41,43,59,60^ and at the neuro-muscular junction^61,62^. Although their expression in neurons is documented, their presence in CG or NSCs had not been reported, prompting us to investigate their localization in the larval CNS.

We first assessed their pattern and level of expression using fluorescent *in situ* RNA hybridizations (HCR RNA-FISH) combined with cell population-specific knockdowns and markers. *dpr10* and *dpr6* were both detected in CG cells, at a lower level for *dpr10* (Figure 3H-K), which also showed an expected neuronal expression (Figure S3A). Knock-in GAL4 lines driven by *dpr10* and *dpr6* enhancers^43^ confirmed such expression pattern (Figure S3B). DIP-α RNA was detected in NSCs as well as in neurons (Figure 3L-M and S3C), and reduced after DIP-α knockdown using a driver (*worniu-GAL4, wor>*) expressed in the NSC and perduring in their larval progeny^27^. While the *DIP-α-GAL4* line showed no NSC signal (Supp. Fig. S3D), possibly due to enhancers missing parts of the expression pattern, antibody staining confirmed DIP*-*α presence in NSCs, lost in a DIP-α null mutant^43^ (Figure S3E).

To visualize the proteins themselves, we created knock-in fluorescent fusions through CRISPR/Cas9 engineering, with either mKate2 in N-terminal or GFP in C-terminal. Strikingly, mKate2∷Dpr10 exhibited a dotted mobile signal travelling along the CG membrane and cytoplasm (Figure 3N and Movie S4). Dpr10::GFP showed a similar speckled expression in the CG along with a faint, continuous membrane localization, and some stronger expression in neurons (Figures S3F-G). The speckled signals of mKate2::Dpr10 and Dpr10::GFP colocalized, while the membrane signal was unique to Dpr10::GFP and the only one present in neurons (Figure S3H and Movie S5). Fluorescent tag localization thus affects Dpr10 dynamics, potentially because C-terminal tagging hinders GPI anchoring. DIP-α::GFP itself combined a dotted pattern and a more continuous signal, both weak except in neurons, and colocalized with a NSC membrane marker (Figure 3O and Movie S6). We wondered whether the two IgSF partners could localize next to each other. Combining mKate2∷Dpr10 with DIP-α∷GFP revealed some dot-level colocalization, sometimes in a transient manner (Figure 3P and Movie S7), suggesting they may function in nearby locations in the CG and NSC, respectively.

To conclude, these different approaches show that Dpr10 and Dpr6 are expressed in the CG while their partner DIP-α is expressed in NSCs. Their dynamic behavior at the protein level may rely on GPI anchoring, suggesting distinct localization patterns based on their specific roles in different cell types.

### DIP-α regulates neural stem cell nuclear shape and proliferation in a cell-autonomous manner

We wondered whether DIP-α in NSC is the functional partner of Dpr10 in CG regulating NSC nuclear shape and proliferation, and whether Dpr6 also plays a role in this process.

First, knocking down *DIP-α* in NSC lineages strikingly recapitulated *dpr10* knockdown in CG, including deformed nuclei (Figures 4A-B), decreased mitotic index and changes in phase distribution towards increased prometaphase/metaphase (Figures 4C-D) in NSCs, while the CG grid-type organization seemed conserved. Similarly to *dpr10* knockdown in the CG, the strength of the mitotic phenotype was variable, with some replicates showing higher mitotic index with either increased prophase or prometaphase/metaphase (Figures S4A-D). Conversely, knocking down *DIP-α* in CG did not result in any obvious phenotype (Figure S4E). Since *wor-GAL4* persists in NSC neuronal progeny^27^, where it drives *DIP-α* knockdown (Figure 3L), we examined whether *DIP-α* loss in neurons contributes to the NSC phenotype. Knocking down *DIP-α* using the neuronal driver *ElaV-GAL4* did not lead to any qualitative change in NSC nuclear shape nor quantitative changes in NSC mitotic index and phase distribution (Figures S4F-H). Further, analysing a *DIP-α* null mutant (*DIP-α^null2^*)^43^ revealed a clear decrease in NSC mitotic index without changes in phase distribution (Figure S4I-K), thus confirming a role of *DIP-α* in regulating NSC proliferation. The overall NSC phenotype, including nuclear shape, was however weaker than for *DIP-α* knockdown in NSCs, suggesting a potential compensation in this constitutive mutant. Of note, *DIP-α::GFP/DIP-α^null2^* flies behaved similarly to *DIP-α^null2^/+* flies regarding NSC proliferation, supporting that this CRISP fusion is functional, at least for this aspect (Figure S4I-K). Taken together, these findings support a cell-autonomous role for *DIP-α* in NSCs, regulating NSC proliferation similarly to how *dpr10* functions in CG. Interestingly, we found that *dpr6* knockdown in CG parallels the effects of *dpr10* knockdown in the CG regarding NSC nuclear shapes, mitotic index and phase distribution, with no detectable change in the CG organization (Figure 4E-H). Of note, while *dpr10* knockdown in NSCs caused no detectable changes in their nuclear shape, *dpr6* knockdown led to mild deformations (Figure S5A). This may be due to cis-binding, indirect regulation, or trans-binding to other DIP partners. The role of *dpr6* in NSCs will not be explored further here.

**Figure 4.**
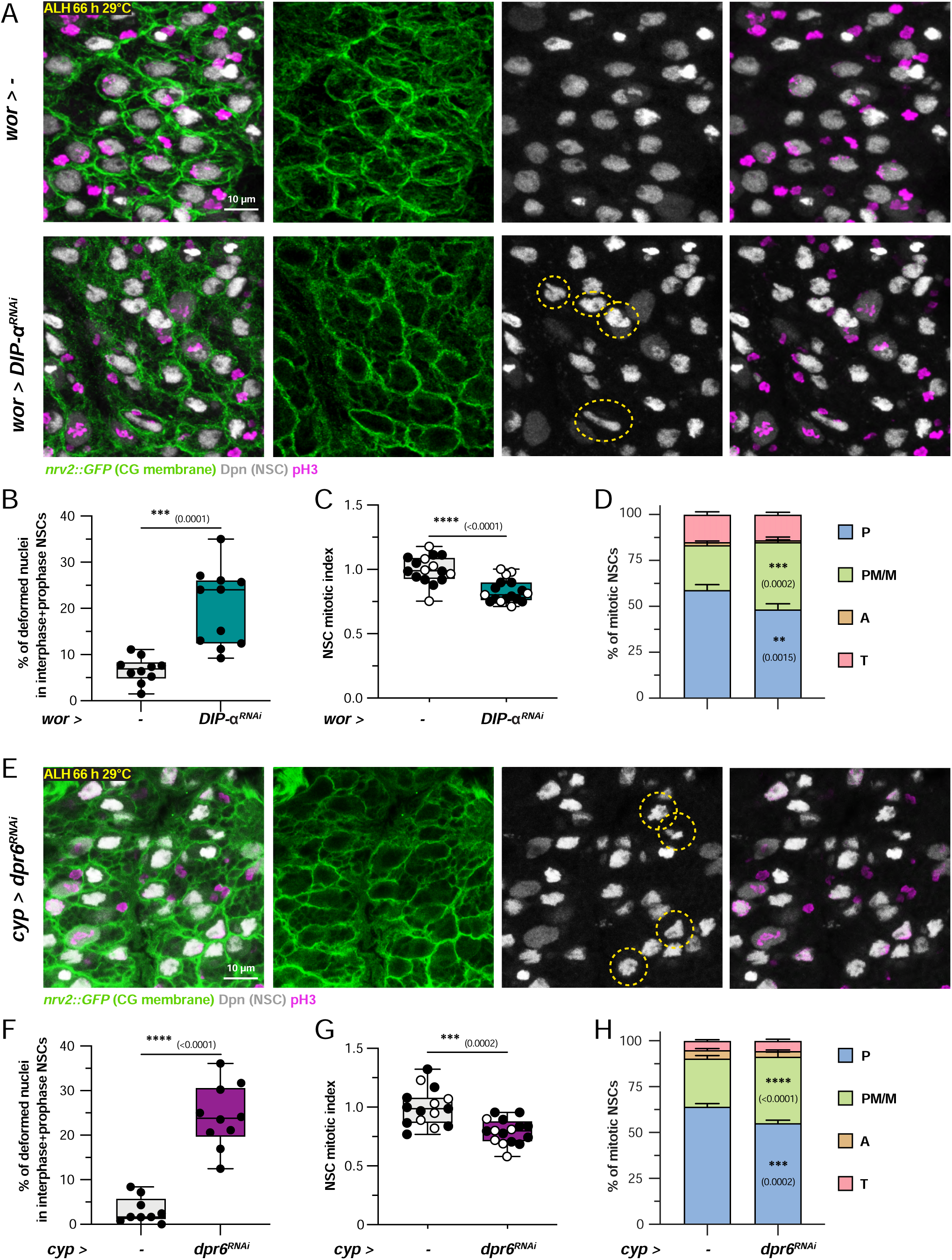
DIP-α is the functional partner of Dpr10 in neural stem cells and can also act trough Dpr6 in the cortex glia, forming an IgSF-CAM^D3^ triad of interactors in the neurogenic niche. (A) Confocal images of the tVNC showing CG membrane and NSC division in control condition (*wor > -*) and under *> DIP-α* knockdown in the CG (*wor > DIP-α RNAi^VDRC104044^)*. Dashed yellow circles indicate examples of deformed nuclei. (B) Box plot of the percentage of NSCs (in interphase + early prophase) showing deformed nuclear shapes in *wor > -* (n = 10 tVNCs) and *wor > DIP-α RNAi^VDRC104044^*(n = 11 tVNCs). Unpaired Student t-test. (C-D) Box plot of mitotic index (normalized to control) and stacked bar chart of phase distribution in NSCs for *wor > -* (n = 16 tVNCs) and *wor > DIP-α RNAi^VDRC104044^* (n = 17 tVNCs). Unpaired Student t-test (C) and two-way ANOVA with Šídák multiple comparison test (D). (E) Confocal images of the tVNC showing CG membrane and NSC division in control condition (*cyp > -*) and under *dpr6* knockdown in the CG (*cyp > dpr6 RNAi^VDRC41161^*). Dashed yellow circles indicate examples of deformed nuclei. (F) Percentage of NSCs (in interphase + early prophase) showing deformed nuclear shapes in *cyp > -* (n = 9 tVNCs) and *cyp > dpr6 RNAi^VDRC41161^*(n = 10 tVNCs). Unpaired Student t-test. (G-H) Mitotic index (normalized to control) and phase distribution in NSCs for *cyp > -* (n = 15 tVNCs) and *cyp > dpr6 RNAi^VDRC41161^*(n = 16 tVNCs). Unpaired Student t-test (G) and two-way ANOVA with Šídák multiple comparison test (H). For box plots, individual values are superimposed and colour-coded to represent multiple experimental replicates. ns, p≥0.05; *, p < 0.05; **, p <0.01; ***, p<0.001; ****, p<0.0001 P: Prophase, PM/M: Prometaphase/metaphase, A: Anaphase, T: Telophase. See also Table S2 and Figures S4-S5.

Single knockdowns in CG suggest that *dpr6* and *dpr10* may not be redundant, but potentially that an equilibrium exists between their complexes, influencing their interactions. Nevertheless, double *dpr10* and *dpr6* knockdowns in the CG produced a similar phenotype regarding nuclear shape and mitotic parameters (Figure S5B-H). This result does not allow us to support a complete non-redundancy of *dpr10* and *dpr6*.

Taken together, these data show that DIP-α is required autonomously in NSCs and its partners Dpr10 and Dpr6 are required non-autonomously in CG for controlling NSC nuclear shape and division. Combined with the known bindings between these IgSF-CAMs, their role in CG-NSC membrane proximity (Figure 3B-D), and their expression profiles and protein localizations in NSCs and CG (Figures 3H-P), we propose that DIP-α establishes heterophilic interactions with its partners Dpr10 and Dpr6 across NSC-CG membranes, which regulate NSC proliferation and nuclear shape.

From now on, we will use either the knockdown of *DIP-α* in NSCs or the knockdown of *dpr10* and/or *dpr6* in CG to disrupt the interactions within this triad, a condition we will call IgSF-CAM^D3^ disruption.

### IgSF-CAM^D3^ interactions regulate tensile forces in the cortex glia, which in turn control nuclear shape and proliferation of the neural stem cells

Our findings have connected disrupted interactions between the CG and NSC membranes to altered NSC functions. What is the nature of the signal conveyed between these two cell types and transduced from one cellular compartment to the other?

Adhesion molecules link cells mechanically across the extracellular space and can connect intracellularly to the cytoskeleton, behaving as mechanotransducers that propagate structural and mechanical changes^63–65^. Cell-cell adhesions not only transmit but also generate mechanical cues^66^. Intriguingly, our observations suggest an alteration in mechanical forces due to IgSF-CAM^D3^ disruption. First, complex changes in nuclear shapes have been recorded as a consequence of uncompensated nuclear deformation following cell mechanical stress^67,68^. Second, cell geometry is a well-known product of intrinsic cell mechanics and extrinsic forces^69^. We tested the hypothesis that IgSF-CAM^D3^ disruption changes the mechanical environment of the NSCs, generating altered forces responsible for NSC nuclear and proliferative phenotypes. This would imply that the CG provide mechanical cues to the NSCs.

The tensile force at the cell cortex is one major contributor to a cell’s mechanical state and has been extensively associated to adhesion and cytoskeletal complexes, notably to the actomyosin network, a major scaffolding yet mechanically responsive component made by filamentous actin and Myosin II^70,71^. Myosin II contractile activity is regulated by its phosphorylation by kinases like RhoA and ROCK, which results in increased cortical tension^72–74^.

We asked whether IgSF-CAM^D3^ disruption affected cortical tension in the NSC niche. Cortical tension can be estimated in live samples by performing targeted laser ablation on the cell membrane and measuring the subsequent recoil velocity of the intact membrane outside of the wound, which reflects its tension magnitude prior to ablation^75,76^. We performed laser ablation on combined *dpr6* and *dpr10* knockdowns in CG, with membranes labeled by Nrv2::GFP, and compared them to controls. In addition, CG-specific expression of a constitutively active Rok (Rok^CA^), the fly ortholog of ROCK, was used as a positive control. We tracked the displacement of landmarks in the CG membrane on each side of the laser cut and measured the initial displacement velocity (Figures 5A-C and Movies S8-S10). We observed a rise in average initial displacement velocity between control (13.71 μm.s^-1^) and Rok^CA^ overexpression (57.75 μm.s^-1^), confirming that Rok^CA^ increased CG tension, which was detectable in our setup. Strikingly, the average initial displacement velocity under *dpr6* and *dpr10* double knockdown (58.28 μm.s^-1^) was similar to that of Rok^CA^ overexpression. These results show that cortical tension is increased in the CG upon IgSF-CAM^D3^ disruption.

**Figure 5.**
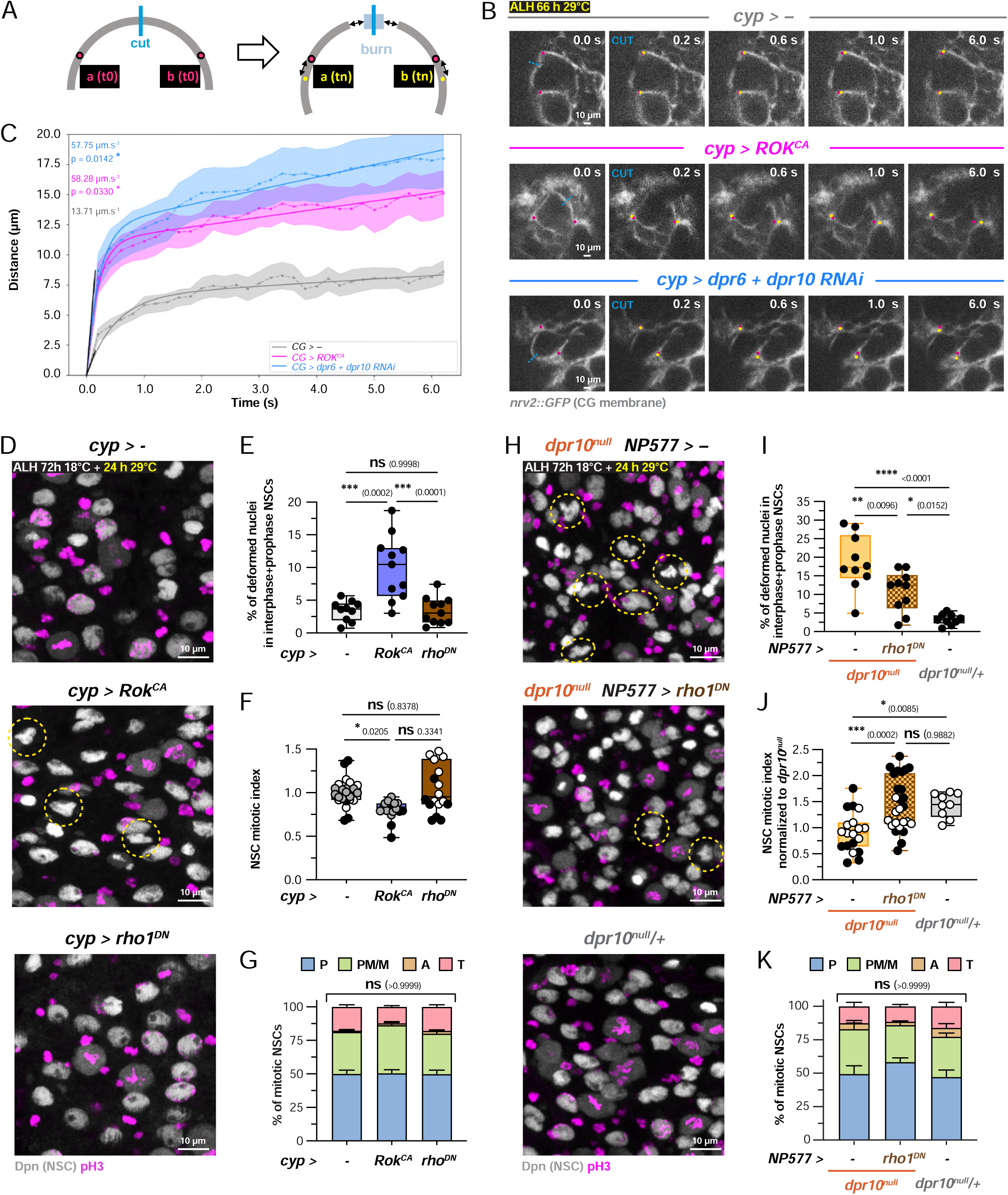
IgSF-CAM^D3^-dependent regulation of cortical tension in the cortex glia is required to maintain normal neural stem cell proliferation and nuclear shape. (A) Schematic of the laser ablation protocol to assess cortical tension pre-ablation. The displacement of two cortical landmarks a and b will be followed from t(0) (magenta dots) until t(n) (yellow dots) to determine initial speed at t(0). (B) Still images of a time-lapse movie picturing the displacement of the CG cortex (grey) after laser ablation, following the protocol shown in A) for control (*cyp > -*, Movie S8), activation of actomyosin contractility in the CG (*cyp > ROK^CA^*, Movie S9) and knockdown of both *dpr10* and *dpr6* in the CG (*cyp>dpr6 RNAi^VDRC41161^+dpr10 RNAi^BDSC27991^*, Movie S10). Dashed cyan lines represent the place of cut. (C) CG membrane displacement versus time graphs for control (n = 15 movies), activation of actomyosin contractility in the CG (n = 15 movies) and knockdown of both dpr10 and dpr6 in the CG (n = 19 movies). Solid lines indicate the mean of the fitted curves while dashed lines indicate the mean of the raw displacement data. Shaded hues represent the SEM of the raw displacement data. d(t0) is represented by lines cutting at the origin and give initial recoil for each ablation. Kruskal–Wallis H test with Dunn’s multiple comparisons test on initial recoil values. (D) Confocal images of the tVNC showing CG membrane and NSC division in control condition (*cyp > -*), constitutive activation of actomyosin contractility (*cyp > ROK^CA^*) and constitutive inhibition of actomyosin contractility (*cyp > rho1^DN^*,). Dashed yellow circles indicate examples of deformed nuclei. (E) Box plot of the percentage of NSCs (in interphase + early prophase) showing deformed nuclear shapes in *cyp > -* (n = 10 tVNCs), *cyp > ROK^CA^* (n = 11 tVNCs) and *cyp > rho1^DN^* (n = 11 tVNCs) from D). One-way ANOVA. (F-G) Blox plot of mitotic index (normalized to control condition *cyp > -*) and stacked bar chart of phase distribution in NSCs for *cyp > -* (n = 25 tVNCs), *cyp > ROK^CA^*(n = 17 tVNCs) and *cyp > rho1^DN^* (n = 18 tVNCs) from D). Nested one-way ANOVA (F) and two-way ANOVA with Šídák multiple comparison test (G). (H) Confocal images of the tVNC showing CG membrane and NSC division in homozygous *dpr10^null^* mutants, homozygous *dpr10^null^* mutants plus transient (24 h at 29°C) inhibition of actomyosin contractility in the CG (NP577 > Rho^DN^), using the *NP577* GAL4 line driving in the CG, and heterozygous *dpr10^null^*mutants. Dashed yellow circles indicate examples of deformed nuclei. (I) Box plot of the percentage of NSCs (in interphase + early prophase) showing deformed nuclear shapes in homozygous *dpr10^null^*mutants (n = 10 tVNCs), homozygous *dpr10^null^* mutants plus transient Rho^DN^ expression in the CG (n = 10 tVNCs) and heterozygous *dpr10^null^* mutants (n = 9 tVNCs) from N). One-way ANOVA. (J-K) Box plot of mitotic index (normalized to control condition a) and stacked bar chart of phase distribution in NSCs for homozygous *dpr10^null^* mutants (n = 18 tVNCs), homozygous *dpr10^null^* mutants plus transient Rho^DN^ expression in the CG (n = 26 tVNCs) and heterozygous *dpr10^null^* mutants (n = 9 tVNCs) from D). One-way ANOVA (J) and two-way ANOVA with Šídák multiple comparison test (K). The yellow font indicates the timing of Rok^CA^/Rho1^DN^ induction. For box plots, individual values are superimposed and colour-coded to represent multiple experimental replicates. ns, p≥0.05; *, p < 0.05; **, p <0.01; ***, p<0.001; ****, p<0.0001 P: Prophase, PM/M: Prometaphase/metaphase, A: Anaphase, T: Telophase. See also Table S2 and Figure S6.

We next investigated whether such increase was functionally relevant for the nuclear and proliferative phenotypes of the NSCs, and the level of interplay between cortical tension in the CG and NSC behaviour. We first assessed the impact of modulating CG cortical tension on NSCs, and especially whether its increase would mimic the phenotype seen under IgSF-CAM^D3^ disruption. We expressed Rok^CA^ or Rho1^DN^, a dominant-negative form of the fly ortholog of RhoA, in the CG to respectively increase or decrease cortical tension in these cells. We limited their overexpression to the mid-larval stage (24 hours at 29°C) to avoid potential effects during earlier NSC reactivation. Expressing Rok^CA^ in the CG resulted in moderately deformed NSC nuclei (Figure 5D-E). Additionally, we observed a reduced mitotic index, with a slight shift towards prometaphase/metaphase (Figure 5F-G), as well as some variability in phenotypic strength (Figures S6A-C), both consistent with the effects seen in *dpr10/dpr6* and *DIP-α* knockdowns. In contrast, expressing Rho1^DN^ in the CG caused no significant changes in NSC nuclear shape, mitotic index, or phase distribution (Figures 5D-G). We conclude that increased CG cortical tension impacts NSC behaviour, including nuclear and mitotic features, in a similar fashion to IgSF-CAM^D3^ disruption.

We then tested whether decreasing cortical tension in CG would be sufficient to rescue the NSC phenotype associated with IgSF-CAM^D3^ disruption. We used a *dpr10^null^* subviable homozygous mutant^43^, which displayed altered nuclear shape and mitotic features compared to a control heterozygous condition (*dpr10^null^/+*) (Figures 5H-K). Expressing Rho1^DN^ specifically in the CG and in a limited fashion (24 h at mid-larval stage) reduced the severity of nuclear deformation, assessed quantitatively by a lower occurrence of abnormal shapes and qualitatively by milder deformations. Additionally, Rho^DN^ expression significantly improved the mitotic index, bringing the mean closer to control levels, though with a wider distribution. The impact on mitotic phases was difficult to assess, as none of the three conditions displayed significant differences from one another. These results indicate that the nuclear and proliferative NSC phenotypes caused by IgSF-CAM^D3^ disruption depend on the increase in CG tensile forces.

Taken together, our findings demonstrate that the control of CG cortical tension by IgSF-CAM^D3^ interactions is required for proper NSC proliferation and regulates NSC nuclear shape.

### Neural stem cells exhibit abnormal mitotic spindles and DNA under IgSF-CAM^D3^ disruption

We then sought to investigate the molecular and cellular mechanisms in NSCs which are influenced by CG mechanical state, and which drive their proliferative and nuclear phenotypes. Alterations in mitotic phase distribution and the live-imaging data suggested that mitosis-linked processes starting from prophase and essential during prometaphase/metaphase were affected by IgSF-CAM^D3^ disruption.

We thus stained for α-tubulin and centrosomin (cnn), a component of the pericentriolar material recruited during centrosome maturation^77,78^, to monitor the mitotic spindle in NSCs. Under *DIP-α* knockdown in NSCs, most prometaphase/metaphase NSCs displayed two centrosomes with a separation distance comparable to control (Figures S7A-B) and a fibre-like central spindle. Spindle morphology however showed striking disruptions, classified qualitatively and quantified across all prometaphase/metaphase NSCs (Figure 6A-B). In control condition, we mostly recorded normal (long and tightly packed central fibre) and diffuse (slightly larger and less long central fibre) morphologies. Under IgSF-CAM^D3^ disruption however, four main other morphology classes were observed: i) large (> 4 μm), often displaying loose packing; ii) bent, in which the spindle appeared curved; iii) misanchored, in which the mitotic spindle is not axial to the centrosomes; and iv) multipolar, in which spindle bipolarity is lost and several, shorter spindles segregate multiple DNA pools. Quantifying spindle width confirmed its shift towards larger and looser morphologies under IgSF-CAM^D3^ disruption (Figure 6C). This resulted in a smaller ratio between DNA and spindle widths (Figure 6D) while the former appears similar to control (Figure S7C), suggesting incorrect chromosomal attachment, a known consequence of spindle defects^79^. Accordingly, metaphase DNA under IgSF-CAM^D3^ disruption displayed a range of aberrant morphologies (Figure 6E), including not centered at the equator, misoriented (not strictly perpendicular to the spindle) and misaligned, a situation associated with improper chromatid segregation^79^. These changes were quantified as a decrease in DNA roundness (Figure 6F), illustrating a more contracted, indented DNA shape. In *Drosophila* NSCs, cortical polarity ensures correct spindle orientation for asymmetric division^80,81^. We stained for α-tubulin along with the asymmetric determinant aPKC, whose is strongly polarized cortically during mitosis^82^ and found correct alignment between aPKC localization and the mitotic spindle’s axis under DIP-α knockdown (Figures S7D-E). Taken together, these data show that IgSF-CAM^D3^ interactions control spindle morphology, but not polarization, in NSCs.

**Figure 6.**
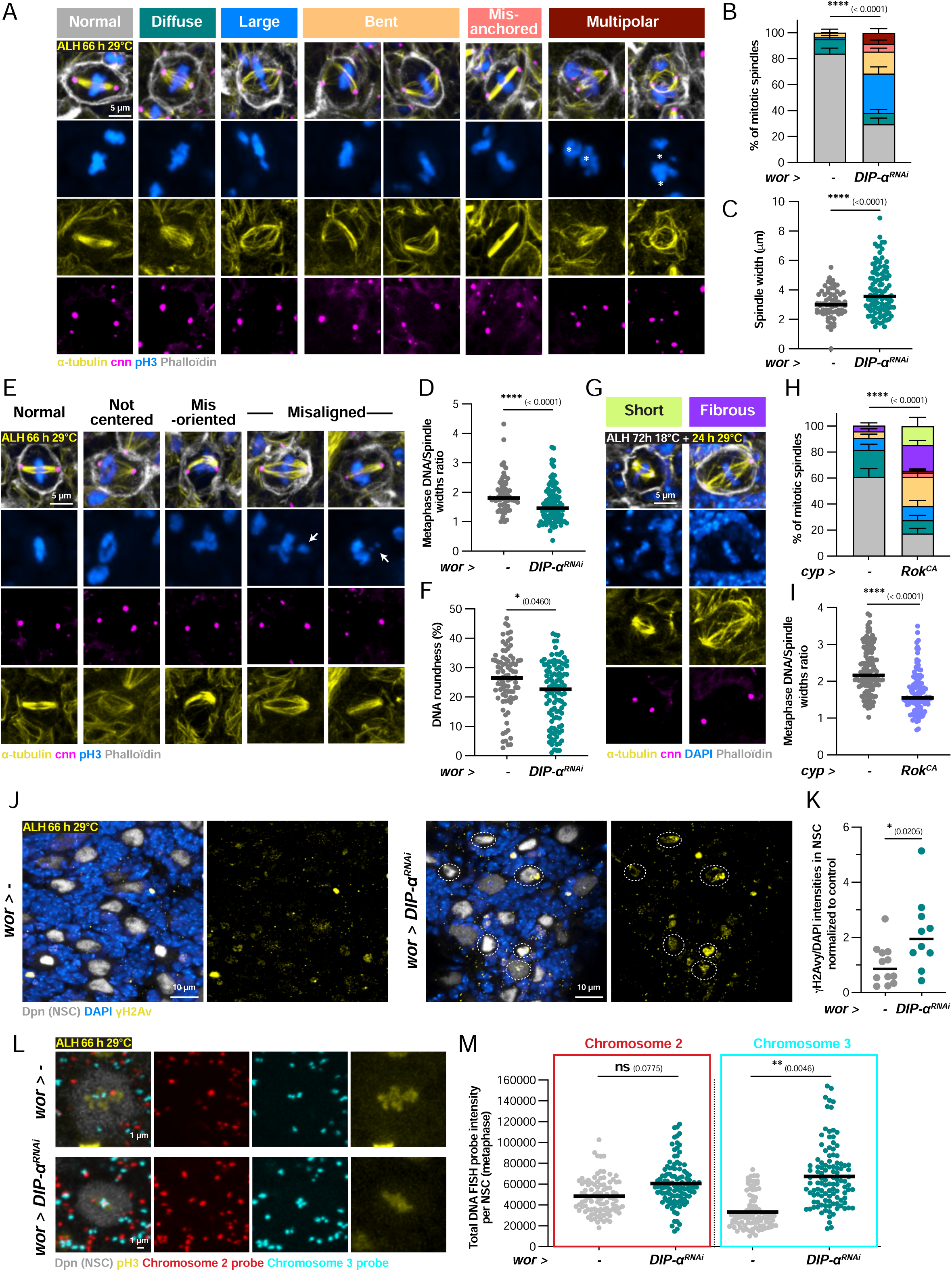
IgSF-CAM^D3^ disruption results in abnormal mitotic spindles and DNA segregation together with DNA damage. (A) Main classes of mitotic spindle morphology (detected by α-tubulin and cnn stainings) found in control and *DIP-α* RNAi in NSCs. Asterisks indicate different genomic copies within one NSC. (B) Stacked bar chart of the proportion of morphological classes from (A) in *wor* > - (n = 7 tVNCs and 67 NSCs) and *wor > DIP-α RNAi^VDRC104044^* (n = 9 tVNCs and 105 NSCs). Two-way ANOVA. (C) Dot plot of spindle length width in *wor* > - (n = 7 tVNCs and 67 NSCs) and *wor > DIP-α RNAi^VDRC104044^* (n = 9 tVNCs and 105 NSCs). Dot = NSC. Mann-Whitney U test on average values per tVNC. (D) Range of metaphase DNA alignment defects found in control and *DIP-α* RNAi in NSCs. Arrows indicate not properly aligned, lagging DNA. (E-F) Dot plots of the ratio between DNA width and spindle width and of DNA roundness for *wor* > - (n = 7 tVNCs and 67 NSCs) and *wor > DIP-α RNAi^VDRC104044^* (n = 9 tVNCs and 105 NSCs). Dot = NSC. Mann-Whitney U test (E) and Nested t-test (F) on average values per tVNC. (G) Additional classes of spindle morphology seen under transient increase of tension in the CG (*cyp > ROK^CA^*, induction for 24 h at 29°C at early L3 stage). (H) Stacked bar chart of the proportion of morphological classes in *cyp > -* (n = 10 tVNCs and 117 NSCs) and *cyp > ROK^CA^*(n = 10 tVNCs and 112 NSCs) from G). Two-way ANOVA. (I) Dot plot of the ratio between DNA and spindle widths for *cyp > -* (n = 10 tVNCs and 117 NSCs) and *cyp > ROK^CA^* (n = 10 tVNCs and 112 NSCs) from G). Dot = NSC. Nested t-test on average values per tVNC. (J) Confocal images of NSCs stained for the DNA damage marker γH2Av (yellow) in control (*wor* > -) and DIP-*α* knockdown (*wor > DIP-α RNAi^VDRC104044^*) in NSCs. Dashed white ovals highlight examples of NSC with high γH2Av staining. (K) Dot plot of γH2Av intensity per NSC (normalized to DAPI intensity and then to the control condition) in *wor* > - (n = 12 tVNCs and 2023 NSCs) and *wor > DIP-α RNAi^VDRC104044^* (n = 10 tVNCs and 1706 NSCs). Dot = tVNC. Unpaired Student t-test. (L) Confocal images of metaphase NSCs stained for oligonucleotide probes (DNA FISH) for chromosomes 2 (red) and 3 (cyan) for control (*wor* > -) and DIP-*α* knockdown in NSCs (*wor* > *DIP-α RNAi^VDRC104044^*). (M) Dot plots of total probe intensity per NSC for each chromosome in *wor* > - (n= 8 tVNCs and 95 NSCs) and *wor > DIP-α RNAi^VDRC104044^*) (n = 9 tVNCs and 116 NSCs). Dot = NSC. Nested t-tests on average values per tVNC. The yellow font indicates the timing of induction. ns, p≥0.05; *, p < 0.05; **, p <0.01; ***, p<0.001; ****, p<0.0001 See also Table S2 and Figure S7.

We then tested whether increased cortical tension in the CG was sufficient to induce defective spindle morphology. Spindle’s analysis under Rok^CA^ overexpression in the CG revealed several abnormalities, including larger, bent, and misanchored structures (Figures 6G-H), similar to those observed in *DIP-α* knockdown in NSCs. A striking case showed one spindle pole not anchored to the centrosome (Figure S7G). Additionally, we found two other spindle categories, short (spindle length < 6 μm, with an extreme case in Figure S7G) and fibrous, where the spindle appeared disorganized with random microtubule fibers instead of a single bundle. The ratio between DNA and spindle widths was also affected (Figure 6I). Increased cortical tension in the CG thus leads to abnormal spindle morphology, producing overlapping phenotypes with the ones observed in IgSF-CAM^D3^ disruption in NSCs. Altogether, these results show that spindle morphology and metaphase DNA assembly are defective under IgSF-CAM^D3^ disruption and depend on CG tensile state. Spindle malformation and improper chromosomal attachments are well associated with prolonged mitotic arrests^83^, thus fitting our earlier observations.

Impairing mitotic processes has been long shown to result in DNA damage and aneuploidy, leading to genomic instability^84^. Prolonged mitotic arrest causes DNA damages^85^ and can be followed by mitotic slippage, when a cell exits mitosis without proper chromosome segregation and cytokinesis^86^, further resulting in aneuploidy. In return, high levels of DNA damage can prevent cells to (re-)enter mitosis, through the activation of the DNA damage checkpoint^87,88^.

We first assessed DNA damage in NSCs, staining for γH2Av, a double DNA strand breaks marker (Figures 6J-K and S7H-K)^89–91^. γH2Av levels appeared increased in the NSC population both under *DIP-α* knockdown in NSCs and *dpr10* knockdown in the CG compared to controls, with some NSCs showing a strong signal. Next, we performed DNA FISH on chromosomes 2 and 3 (two out of the four *Drosophila* chromosomes) (Figure 6L) and measured the total intensity of each probe per NSC as an indicator of DNA content, focusing on metaphase to ensure consistent replicative status. While chromosome 2 probe displayed a restricted, non-significant increase in total intensity, chromosome 3 probe showed a doubling (Figure 6M). This result suggests that DNA content changes occur in NSCs under IgSF-CAM^D3^ disruption, leading to aneuploidy at certain genetic loci. The discrepancy between probes could be explained by known biases in the frequency of aneuploidy between DNA regions^92^.

Altogether these data show that IgSF-CAM^D3^ disruption leads to malformation of the mitotic spindle, further resulting in improper DNA positioning and segregation during mitosis, with DNA damage and some level of aneuploidy.

### Nuclear lamina shape and composition are altered under IgSF-CAM^D3^ disruption

The nuclear lamina is a dense, fibrous network primarily composed of intermediate filament proteins called lamins (A/C- and B-types) and their associated proteins^93,94^. It is key in controlling the shape and mechanical rigidity and stability of the nucleus under a range of cell processes^95^.

*Drosophila* has one B-type (Dm0 or Dmel/Lam) and one A-type (Dmel/LamC) lamin. As reported before^96^, larval NSCs do not stain for LamC (Figure S8A) implying that the B-type lamin Lam (and its partners) will be sole responsible for regulating the lamina-dependent mechanics of the nuclear envelope. Lam staining under *DIP-α* knockdown in NSCs outlined highly contorted shapes of the nuclear lamina, displaying many folds and indents towards the inner nuclear compartment compared to control (Figure 7A). Quantifying the proportion of Lam staining area compared to the whole nuclear area at the equatorial plane confirmed this observation (Figure 7B). Comparing Lam levels in control and *DIP-α* knockdown also revealed that Lam levels were higher in NSCs (Figure 7C). Strikingly, both Lam distorted shapes and higher levels were recapitulated under *Rok^CA^*overexpression in CG (Figures 7D-F). Increased cortical tension in the CG is thus sufficient to deform the nuclear lamina and increase Lamin content. Our findings show that IgSF-CAM^D3^ interactions control the shape and composition of the nuclear lamina in NSCs, which is sensitive to tensile forces in the CG.

**Figure 7.**
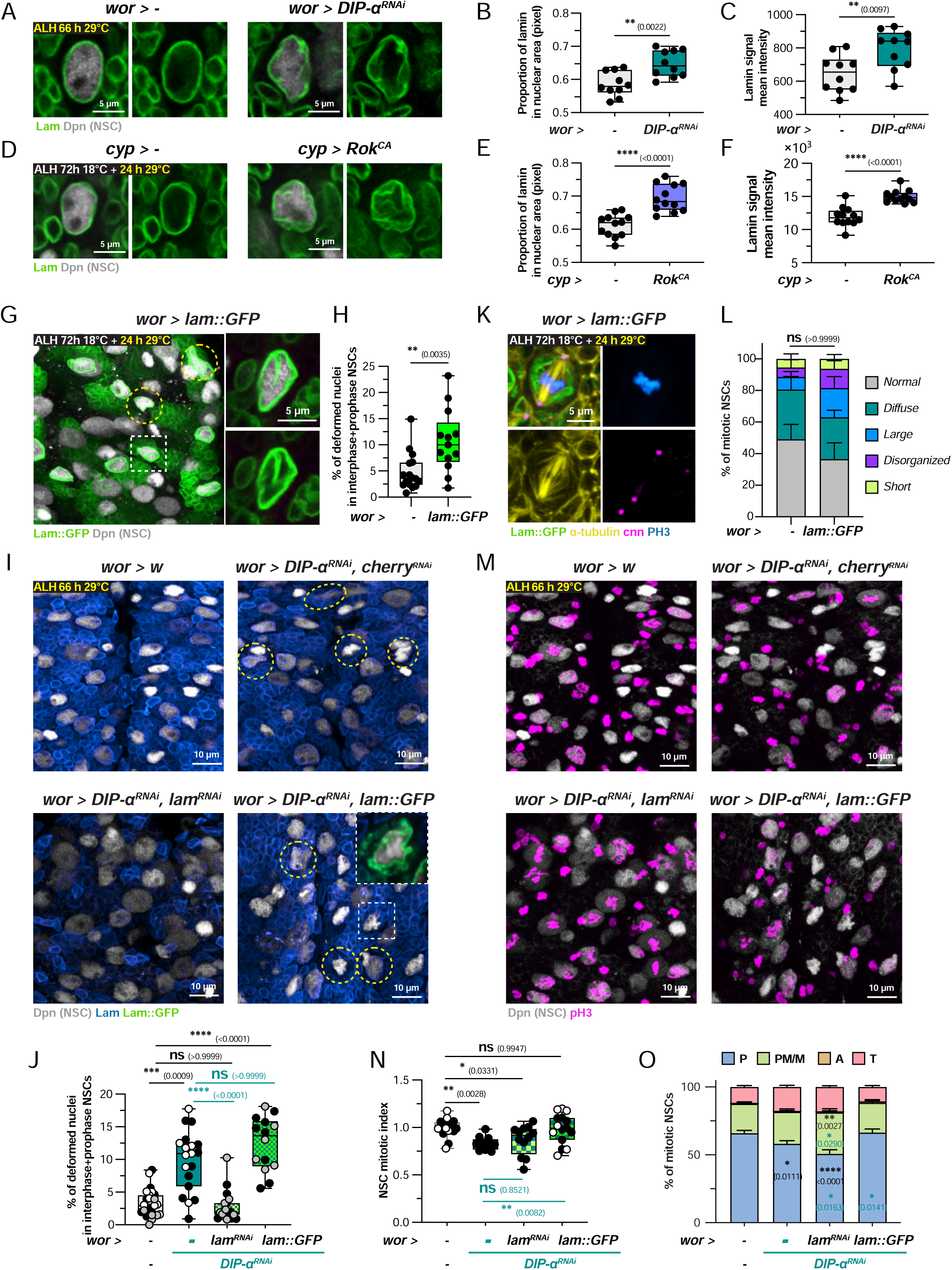
IgSF-CAM^D3^disruption leads to Lamin enrichment in the nuclear lamina of NSCs, causing nuclear deformation while protective of proliferation. (A) Confocal images of the nuclear lamina (anti-Lam, green) in NSCs for control and *DIP-α* RNAi in NSCs. (B-C) Box plots of the proportion of lamin signal in the total nuclear area (in pixel) as a proxy for folding and thickness of the nuclear lamina (B) and of lamin mean intensity (C) in *wor* > - (n = 10 tVNCs and 376 NSCs) and *wor > DIP-α RNAi^VDRC104044^* (n = 10 tVNCs and 375 NSCs). Linear mixed models on average values per tVNC. (D) Confocal images of the nuclear lamina (anti-Lam, green) in NSCs for control (cyp > -) and transient tension increase (24 h) in the CG (*cyp > ROK^CA^*). (E-F) Box plots of the proportion of lamin signal in the total nuclear area (in pixel) (E) and of lamin mean intensity (F) in *cyp > -* 1(n = 12 tVNCs and 327 NSCs) and *cyp > ROK^CA^* (n = 12 tVNCs and 340 NSCs) from D). Linear mixed models on average values per tVNC. (G) Confocal image of a tVNC under Type-B lamin (Lam) overexpression in NSCs (*wor > lamin::GFP*). Lam was expressed for 24 h at 29°C. Dashed yellow circles show examples of deformed nuclei. Close-ups (single plane, at the equatorial) highlights Lam folding. (H) Box plot of the percentage of NSCs (in interphase + early prophase) showing deformed nuclear shapes in *wor > -* (n = 16 tVNCs) and *wor > lamin::GFP* (n = 13 tVNCs). Mann-Whitney U test. (I) Confocal images of the tVNC showing NSC nuclear shape (Dpn, grey) and Lam staining (blue) in control condition (*wor > -*), dose-compensated *DIP-α* knockdown in NSCs (*wor > DIP-α RNAi^VDRC104044^ + mCherry RNAi*), double *DIP-α* and *Lam* knockdown in NSCs (*wor > DIP-α RNAi^VDRC104044^ + lam RNAi*) and *DIP-α* knockdown plus lamin overexpression in NSCs (*wor > DIP-α RNAi^VDRC104044^ + lam::GFP*). Dashed yellow circles indicate examples of deformed nuclei. The insert shows an example of Lam staining in *lam::GFP* overexpression under *DIP-α* knockdown. (J) Box plot of the percentage of NSCs (in interphase + early prophase) showing deformed nuclear shapes in *wor > -* (n = 11 tVNCs), *wor > DIP-α RNAi^VDRC104044^ + mCherry RNAi* (n = 11 tVNCs), *wor > DIP-α RNAi^VDRC104044^ + lam RNAi* (n = 11 tVNCs) and *wor > DIP-α RNAi^VDRC104044^ + lam::GFP* (n = 8 tVNCs). Kruskal–Wallis H test with Dunn’s multiple comparisons test. (K) Spindle morphology (stained with α-tubulin, yellow) under lamin overexpression (green) in NSCs (*wor > lamin::GFP*). Lam was expressed for 24 h at 29°C. (L) Stacked bar chart of the proportion of morphological classes for the central spindle in *wor* > - (n = 11 tVNCs and 94 NSCs) and *wor > lamin::GFP* (n = 12 tVNCs and 79 NSCs). Two-way ANOVA. (M) Confocal images of the tVNC showing NSC division in control condition (*wor > -*), dose-compensated *DIP-α* knockdown in NSCs (*wor > DIP-α RNAi^VDRC104044^ + mCherry RNAi*), double *DIP-α* and *Lam* knockdown in NSCs (*wor > DIP-α RNAi^VDRC104044^ + lam RNAi*) and *DIP-α* knockdown plus lamin overexpression in NSCs (*wor > DIP-α RNAi^VDRC104044^ + lam::GFP*). (N-O) Blox plot of mitotic index (normalized to control *wor > -*) and stacked bar chart of phase distribution in NSCs for *wor > -* (n = 18 (N) and 12 (O) tVNCs), *wor > DIP-α RNAi^VDRC104044^ + mCherry RNAi* (n = 12 tVNCs), *wor > DIP-α RNAi^VDRC104044^ + lam RNAi* (n = 12 tVNCs) and *wor > DIP-α RNAi^VDRC104044^ + lam::GFPs* (n = 15 (N) and 8 (O) tVNCs). Kruskal–Wallis H test with Dunn’s multiple comparisons test (N) and two-way ANOVA with Šídák multiple comparison test (O). The yellow font indicates the timing of induction. For box plots, individual values are superimposed and colour-coded to represent multiple experimental replicates. ns, p≥0.05; *, p < 0.05; **, p <0.01; ***, p<0.001; ****, p<0.0001 P: Prophase, PM/M: Prometaphase/metaphase, A: Anaphase, T: Telophase. See also Table S2 and Figure S8.

Nuclear shape is known to be altered by the pathological overexpression of Type-B lamins^97,98^ and linked to excess surface area of the nuclear lamina ^99^. We wondered whether increased Lam levels were causative of increased nuclear deformations seen with IgSF-CAM^D3^ disruption. First, transiently driving a *lam::GFP* fusion construct in NSCs resulted in altered nuclear shapes, albeit at a moderate level and amplitude (Figures 7G-H). Thus, increased Lam levels is sufficient to alter nuclear shape, although the exact level and time of increase might modulate phenotypic strength. To test the dependency of nuclear deformation on Lam levels under IgSF-CAM^D3^ disruption, we co-expressed *lam* and *DIP-α* RNAis in NSCs, compared to a dose-compensated *DIP-α* RNAi (*DIP-α* + *mcherry* RNAis) and controls (Figure 7I). The efficiency of *lam* RNAi was tested beforehand (Figure S8B-D). We found that *lam* RNAi reduced nuclear deformations caused by *DIP-α* RNAi to control levels (Figure 7J). Of note, we found that *lam::GFP* co-expression tended to amplify the severity of nuclear deformations, although in a non-significant manner regarding penetrance. In summary, these findings show that IgSF-CAM^D3^ disruption results in increased Lam levels, which in turn cause nuclear lamina deformations.

### Nuclear deformation is not the cause of mitotic impairment and instead might serve a protective role

Our investigation found that mechanical forces from the CG, regulated by adhesion complexes across CG and NSC membranes, are transmitted to the NSC where they impact both the nuclear lamina and the mitotic machinery. We wondered whether these two aspects of the NSC phenotype were linked as a result of one another, especially since the spindle remains encapsulated by the nuclear envelope throughout mitosis in *Drosophila* NSCs^100^.

We first asked whether Lam increase could explain the alterations in spindle morphology, by providing some constraint during this semi-closed mitosis. NSCs under *lamin::GFP* overexpression did not show abnormal morphologies nor widths of the mitotic spindle and metaphase DNA (Figures 7K-L and S8E-G). We still observed a mild change in the ratio between these spindle and DNA widths, suggesting that DNA attachment in metaphase might be influenced (Figure S8H). Metaphase DNA roundness was also altered (Figure S8I), opposite to the effect seen with IgSF-CAM^D3^ disruption (increase *vs*. decrease, see Figure 6F). Altogether, we found that increasing Lam levels in NSCs is not sufficient to cause defective mitotic spindle morphology.

We then investigated whether altering Lam levels modulated the NSC proliferative defects caused by IgSF-CAM^D3^ disruption, particularly whether reducing Lam could rescue such phenotype. We co-expressed *lam* and *DIP-α* RNAis in NSCs, compared to a dose-compensated *DIP-α* RNAi (*DIP-α* + *mcherry* RNAis) which showed decreased mitotic index and altered phase distribution compared to control (Figures 7M-O). Co-expressing *lam* RNAi had no significant impact on the mitotic index but exacerbated the shift toward higher prometaphase/metaphase levels associated with *DIP-α* knockdown. This result indicates that reducing Lam levels is insufficient to rescue NSC proliferation and may even exacerbate its impairment. Combined with the *lamin::GFP* overexpression results, we propose that Lamin increase by itself is not causative of the mitotic defects resulting from IgSF-CAM^D3^ disruption. Furthermore, since proliferation is not restored -and may even worsen-when nuclear deformations themselves are fully rescued (see Figure 7J), we propose that nuclear deformation and mitotic defects are not linearly connected in this context.

The worsening of NSC mitotic phenotype caused by IgSF-CAM^D3^ disruption under *lam* RNAi made us curious as whether there still was some influence of Lam levels *per se* on NSC proliferation. Surprisingly, we found that co-expressing *lam::GFP* with *DIP-α* RNAi fully rescued both the changes in mitotic index and phase distribution associated with *DIP-α* knockdown alone (Figures 7M-O). Together with mitotic defects worsening under *lam* RNAi, this result suggests that Lam levels modulate NSC proliferation under IgSF-CAM^D3^ disruption. We also found that Lam levels modulate NSC proliferation in normal conditions, as both lowering and increasing them (through *lam* RNAi and overexpression of a *lam::GFP* respectively) led to increased mitotic index, accompanied by increased prometaphase/metaphase for *lam::GFP* overexpression only (Figures S8J-L). Thus, while Lam-based nuclear deformations are not causative of mitotic defects, the regulation of Lam levels themselves still impacts NSC proliferation. The full rescue of NSC mitotic phenotype by Lam overexpression also suggests that elevated Lam provides some protective mechanism to mitotic processes in NSCs.

## DISCUSSION

Our study shows that niche mechanics are transmitted to and regulate NSC function. We identified a triad of IgSF-CAMs (Dpr10/Dpr6 in CG and DIP-α in NSCs) which physically and mechanically link the NSC and CG membranes while regulating CG tensile state. Their disruption leads to an increase in CG cortical tension, which in turn is both sufficient and necessary to impair mitosis in NSCs. Mitotic defects are marked by spindle abnormalities and delayed mitotic progression, eventually preventing the cell from re-entering mitosis. IgSF-CAM^D3^-mediated increase of CG cortical tension also deforms NSC nuclei, increasing Lamin as a protective response to mechanical stress. Overall, our results highlight niche mechanics as key orchestrators of NSC function and identify IgSF-CAMs as mechanoregulators.

The link between IgSF-CAM adhesions and CG tension raises several questions. First, why does the tensile state of one cell type change when its interaction with another is disrupted? Could NSCs provide feedback that affects CG mechanical states? CG contractility might vary, potentially locally, with NSC cycling and be influenced by the neurogenic output of NSCs. Second, what intracellular mechanisms implement tension changes in CG? While we used actomyosin-based contractility to manipulate CG mechanics, whether it reflects its importance during normal processes remains unclear. We did not observe increased Myosin II levels under IgSF-CAM^D3^ disruption (Figure S2F), suggesting alternative mechanisms or limitations in detecting localized changes in the CG. Microtubules also are both sensors and providers of mechanical forces^101,102^ and it would be worthwhile to test their involvement in CG.

Upstream to these interrogations lies the central question of how DIP-α, Dpr10 and Dpr6 molecularly impinge on mechanical forces at the cell membrane. These IgSF-CAMs typically lack cytoplasmic domains or have very short cytoplasmic regions with no known functional domains or signaling motifs. This supports a model where these function as adhesion and specificity receptors but must rely on co-receptors which possess transmembrane and cytoplasmic regions to relay signals intracellularly^58^. These co-receptors could directly bind cytoskeletal components and the mechanotransduction machinery, with the mechanosensitive channel Piezo appearing as an appealing candidate. The mystery of IgSF-CAMs’ roles in mechanotransduction is reinforced by the cellular behaviour of knock-in fusions, which appears as motile dots along the membranes. Whether they are constantly relocalising, adapting to local membrane changes and following or hooking on their co-receptors, remains to be understood.

An intriguing result is the distinct roles of Dpr10 and Dpr6 in the CG. Knockdown of each alone affects NSC division, suggesting non-redundant functions. Dpr6 and Dpr10 may have similar roles in regulating membrane tension through DIP-α binding but act at different times in the cell cycle. They might also regulate each other’s expression, affecting DIP-α interactions. Variations in affinity could influence how fast cells form or break interactions, helping to remodel local contacts quickly. Their imbalance, rather than a complete disruption, may impact NSC behaviour. Finally, they might act through their other DIP partners, DIP-β and DIP-λ.

NSC response to disrupted cell-cell adhesion and increased CG tension is double. First, nuclear deformation happens alongside elevated Lamin levels. Lamin A/C typically provides nuclear rigidity and scales with tissue stiffness^67,103^, while Lamin B may play this role in its absence^104^. Here, we propose that CG tension-induced increase in the B-type Lam results in nuclear deformation and stiffening in *Drosophila* NSCs. Both nuclear stiffening and softening protect against mechanical forces to minimize DNA damage (reviewed in^105–107^). Mild stress triggers stiffening of the nuclear shell to provide mechanical resistance and softening of nuclear content to dissipate force^108,109^, but prolonged stress can lead to DNA damage despite further adaptive mechanisms. Here, NSCs exhibit both nuclear stiffening and DNA damage, suggesting that the protective mechanisms are insufficient.

Second, NSC proliferation is altered. The formation of abnormal mitotic spindles is one striking contributing factor, though its underlying mechanisms remain unclear. Investigating known spindle processes linked to the cell membrane and cortex would be important. For example, the spindle is mechanically connected to the cell cortex through astral microtubules. These interactions generate pulling forces that control spindle position and orientation, and are mediated by protein complexes like the Gαi–LGN–NuMA–dynein complex, which links microtubule motors to membrane-anchored factors^110^. In *Drosophila* NSCs however, Mud (NuMA homolog) affects spindle positioning but not morphology^53^, questioning the involvement of other partners identified in vertebrates, such as unconventional myosins^110^. Other interactions linking microtubules and actin could also play a role. The *Drosophila* spectraplakin Shot, known to localize at the spindle poles and influence spindle assembly and chromosome segregation, might be a relevant candidate^111^.

Whether the nuclear and mitotic phenotypes are linked remains to be thoroughly assessed. While nuclear deformation induced by elevated Lam levels is not causative of spindle defects (Figures 7K-L) and is not required for mitotic impairment (Figures 7M-O), Lam still modulates NSC mitosis both in normal conditions and under IgSF-CAM^D3^ disruption (Figures S8J-L and 7M-O). This highlights the existence of a nuclear-based regulation of mitosis, whose extent and mechanisms are yet to be determined. Our finding that boosting Lam levels rescues NSC proliferation, and as such is protective of mitotic processes, is unexpected and especially intriguing. One possible explanation is that increased nuclear stiffening provides enhanced protection against DNA damage, thereby enabling mitotic re-entry that would otherwise be blocked by checkpoint mechanisms. Nuclear and mitotic phenotypes could also result from common mechanotransduction mechanisms relaying CG tensile forces to the NSC. Investigating the respective roles of the actin and microtubule cytoskeleton in this transmission would be relevant. The LINC complex, a key player in nuclear mechanotransduction linking the cell membrane to the nuclear lamina^107,112^, is also a prime candidate, especially given its recent involvement in centrosome positioning^113^. Interestingly, an increase in Lamin following mechanical stress has also been shown to protect the cell cycle in the embryonic heart^114^, highlighting conserved core mechanisms.

While matrix stiffness has been shown to influence oligodendrocyte progenitor function throughout the lifespan^15^, the role of cell-based mechanics remained unprobed. Our findings reveal that the *in vivo* mechanics of their cellular microenvironment control NSC proliferative ability. The abundance of adhesions and tight cellular interactions among NSCs, progenitors, and various niche cells^6,10^, within both developing, mature and ageing mammalian niches, suggests a conserved and broad cell-based mechanical regulation of neurogenesis.

## METHODS

### Fly lines and husbandry

*Drosophila melanogaster* lines were raised on standard cornmeal and maintained at 18°C until their use in crosses and experimental procedures. Lines used in this study are listed below:

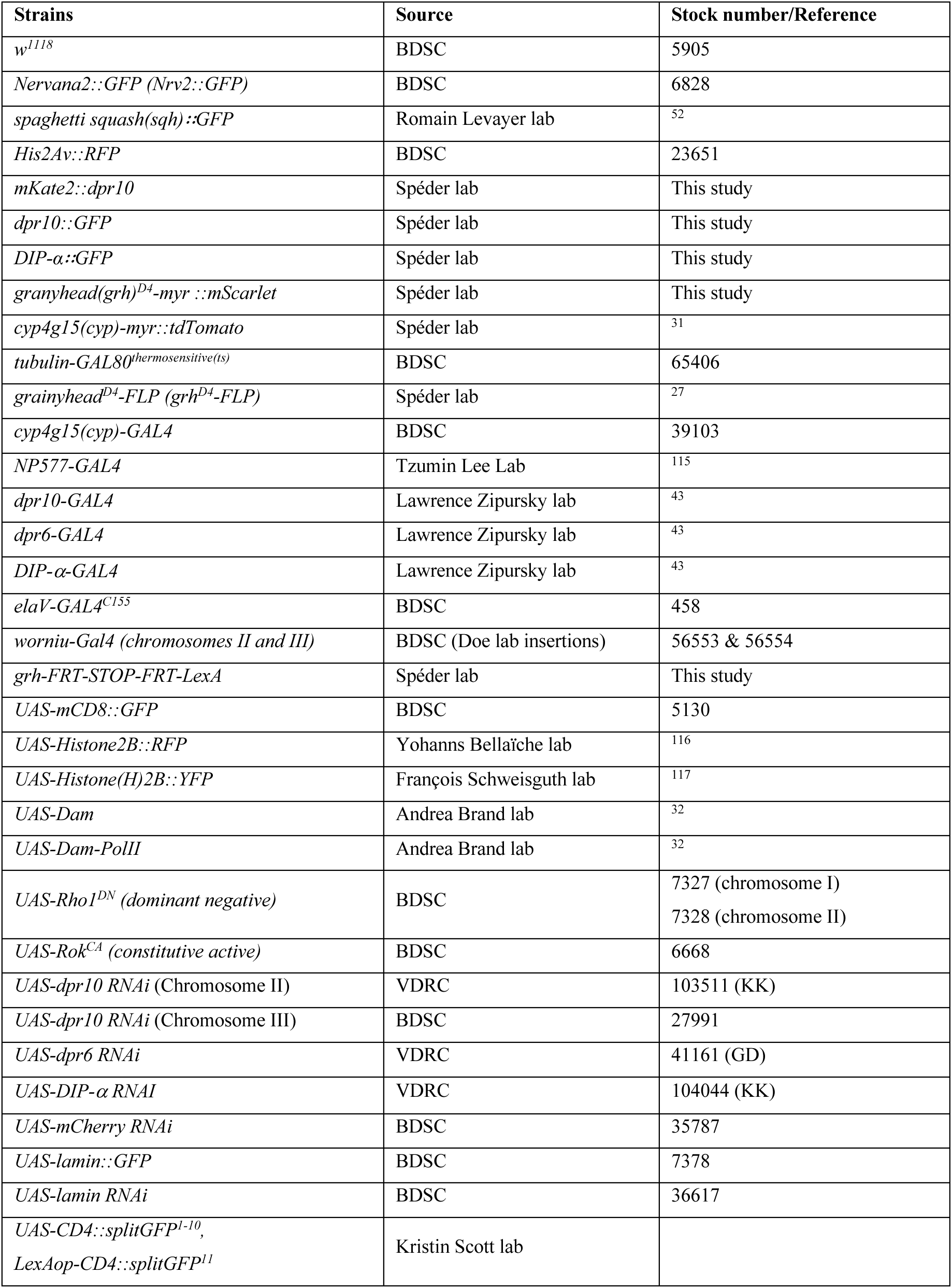
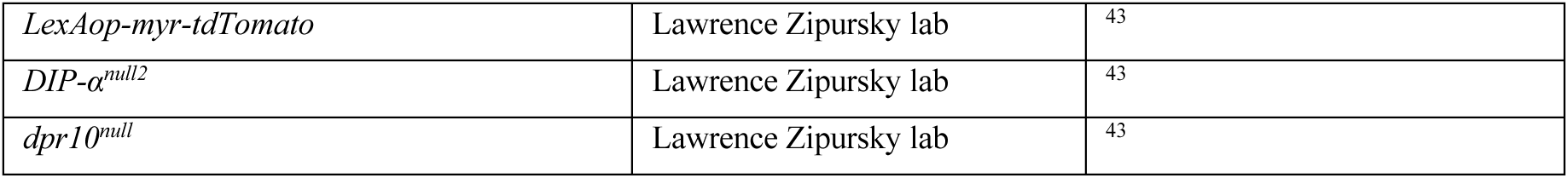

### Larval culture and staging

Embryos were collected within 2-4 hour time window on grape juice-agar plates and kept at either 25°C for 20-24 hours or 18°C for 40-48 h, until the beginning of larval hatching. Freshly hatched larvae were collected within a 1-2 hour time window (defined as 0 hours after larval hatching, ALH0), transferred to fresh yeast paste on a standard cornmeal food plate and raised under chosen experimental conditions (time and temperature) until processing.

25°C was used as the normal developmental temperature for *Drosophila melanogaster*. 18°C was used to keep the thermosensitive allele of GAL80 (GAL80^ts^), a repressor of GAL4, active (so to switch off GAL4 expression), while 29°C was used for its inactivation (so to switch on GAL4 expression). For all experiments using the GAL80^ts^, embryogenesis took place at 18°C, unless noted otherwise. This was to prevent a contribution of embryonic function to the larval phenotype.

Detailed genotypes, crosses and culture regimens are listed in Table S2.

### DNA cloning and *Drosophila* transgenics

The *grainyhead* (*grh*) D4 enhancer (4 kb from the second intron of the *grainyhead* gene), which drives in post-embryonic NSCs^119,120^, was amplified from genomic DNA extracted from *grh (NB)-GAL4* adult flies, with a *hsp70* promoter fused in C-terminal. For creating *grh^D^*^4^*-FRT-STOP-FRT-LexA*, a FRT STOP cassette was amplified from an *UAS-FRT.STOP-Bxb1* plasmid (gift from MK. Mazouni) and the LexA sequence was amplified from the entry vector pENTR L5-LexA::p65-L2 (gift from S. Stowers / M. Landgraf; Addgene 41437). The two amplicons were joined together by overlapping PCRs. This *FRT-STOP-FRT-LexA* amplicon together with the grh^D4_DSCP^ enhancer were inserted in the destination vector pDESThaw sv40 (gift from S. Stowers) using the Multisite gateway system ^121^ to generate a *grh^D^*^4^*-FRT-STOP-FRT-GAL4* construct. The construct was integrated into the fly genome at an attP2 docking site through PhiC31 integrase-mediated transgenesis (BestGene). Several independent transgenic lines were generated and tested, and one was kept.

### Generation of fluorescently tagged CRISPR lines

The CRISPR-associated protein 9 nuclease (Cas9) system was used to insert DNA fragments encoding GFP or mKate2 at the C- or N-terminal of Dpr10 and DIPα generating the following alleles: *DIPα∷GFP*, *dpr10∷GFP* and *mKate2::dpr10*. To direct the Cas9 endonuclease to specific DNA sites, two different and concatenated guide RNAs were cloned into a single *CFD5* plasmid vector under the strong ubiquitous *U6:3* promoter and were preceded by tRNAs ^122^. This strategy uses the endogenous tRNA-processing system to boost the CRISPR/Cas9 system. The oligonucleotides used as guide RNAs for CRISPR/Cas9 target sites are listed below:

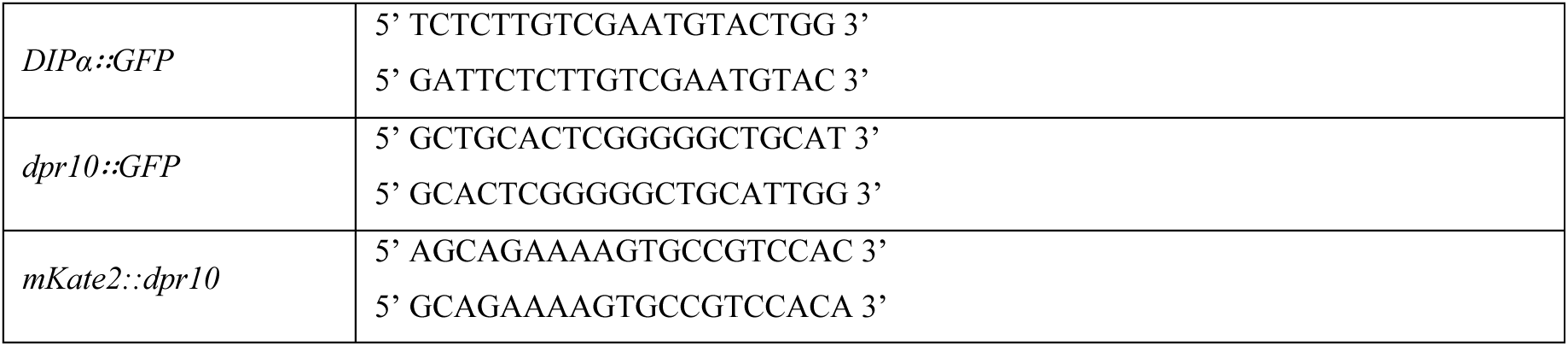

CRISPR/Cas9-mediated homologous recombination was used to insert the fluorescent labels at the specific nucleated sites. Homologous sequences of 1 kb flanking the CRISPR/Cas9 target sites were stacked into plasmid vectors harbouring the C- or N-terminal *GFP* or *mKate2* sequence and a *hsp70∷miniwhite* gene flanked by two loxP sites allowing the removal of the *miniwhite* gene in the final strain by Cre recombinase. The primers used to clone the homologous regions inside the plasmid vector containing the fluorescent tags are indicated below:

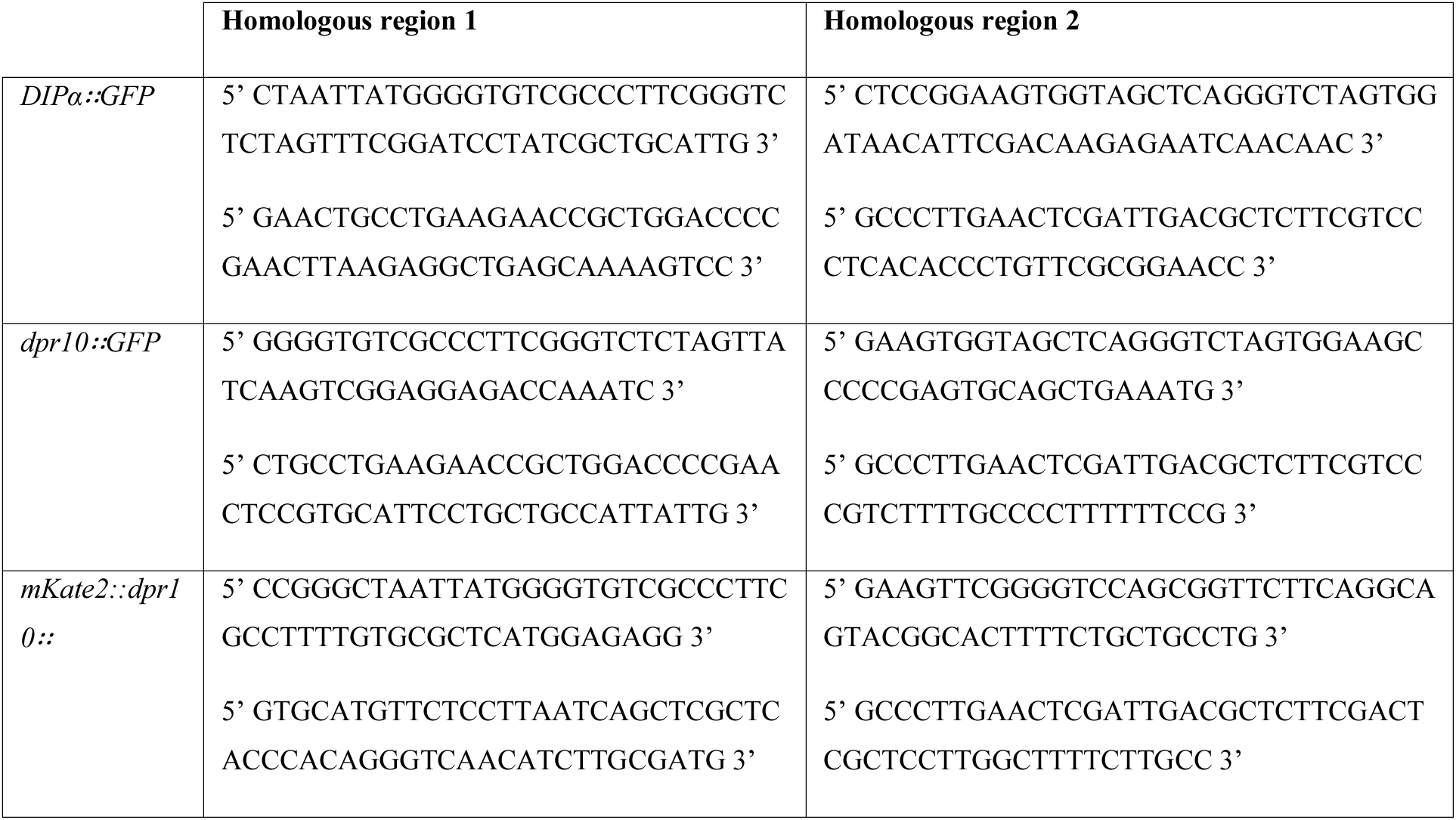

All the cloning was done following the sequence and ligation-independent cloning (SLIC) method ^123^. Both reporter and guide RNA plasmid vectors were injected into *vasa-Cas9* flies (BDSC reference 55821 for DIPα, 51324 for Dpr10) by BestGene®.

### Transcriptional profiling

We used Targeted DamID^32^ to profile transcription in cortex glia at different times along niche formation (T1, T2 and T3). *UAS-Dam* and *UAS-Dam-Pol II* flies were crossed to *tubulin-GAL80^ts^*; *cyp4g15-GAL4* flies. Different tests were performed to center the expression of Dam-Pol II at the right time windows. Each time, progeny was raised at 18°C then shifted to 29°C at the following time: T1: 32 hours after egg laying; T2: 32 hours after larval hatching; T3: 76 hours after larval hatching. In all cases, CNS were dissected out 18 hours after the 29°C shift. Processing of genomic DNA and data analysis were performed as previously described^32–34^. Three biological replicates were performed for each timepoint. We excluded an outlying replicate at T1 that dominated clustering and whose Pearson correlation was < 0.92 to all other replicates (mean Pearson correlation between all other replicates was 0.95). We took log2 Dam ratios (DamPol II occupancy/Dam occupancy) per gene with FDR < 0.01 for all biological replicates, resulting in 3436 genes. To understand relative changes across the time course these were max-normalised and then clustered into 8 groups by k-means clustering.

### Fixed tissue immunohistochemistry

CNS from staged larvae were dissected in PBS and fixed for 20 min in 4% methanol-free formaldehyde (ThermoFisher 28908) diluted in PBS, while rocking at room temperature. They were washed in PBS once and permeabilised in PBS-T (PBS containing 0.3% Triton X-100 (Sigma T9284)) three times for 15 minutes each. CNS were incubated with blocking solution (PBS-T, 5% Bovine Serum Albumin (Sigma A3608), 2% Normal Goat Serum (Abcam ab7481)) for 1 hour at room temperature while rocking. Primary antibody dilution was prepared in blocking solution and CNS were incubated two nights at 4°C, then washed three times for 10-15 minutes each with PBS-T while rocking. Secondary antibodies were diluted in 1:200 in blocking solution and CNS were incubated overnight at 4°C or 3-4 h at room temperature with secondary antibodies. CNS were washed three times in PBS-T while rocking and kept in PBS at 4°C until mounted in homemade Mowiol (Sigma 81381) mounting medium on a borosilicate glass side (number 1.5; VWR International). Primary antibodies used were: guinea pig anti-Dpn (1:5000,^31^), chicken anti-GFP (1:2000, Abcam ab13970), rat anti-ELAV (1:100, 7E8A10-c, DSHB), mouse anti-Repo 1:100 (DSHB, 8D12-c), rabbit anti-Phospho-histone H3 (1:200, Sigma 06570), rabbit anti-Phospho-histone H3 (1:200, Sigma H-0412), mouse anti-Phosphohistone H3 (1:200, Cell Signaling 9706), mouse anti-Lamin Dm0 (1/10, ADL67.10-s, DSHB), mouse anti-Lamin C (1/10, LC28.26, DSHB), rabbit anti-aPKc (1/500, Santa Cruz Biotechnology sc-17781), mouse anti-α-tubulin (1/200, clone DM1A, Cell Signalling Technology #3873), rabbit anti-miranda (1/500; ^124^), guinea pig anti-cnn (1/1000, ^125^), rabbit γH2Av (H2AvD pS137, Rockland 600-401-914), mouse anti-DIP-α (1/10, ^43^). Fluorescently-conjugated secondary antibodies Alexa Fluor 405, Alexa Fluor 488, Alexa Fluor 546 and Alexa Fluor 633 (ThermoFisher Scientific) were used at a 1:200 dilution. DAPI (4′,6-diamidino-2-phenylindole, ThermoFisher Scientific 62247) was used to counterstain the nuclei.

Images were acquired with a confocal microscope (Zeiss LSM880) on a ZEN software, with an optimal distance between each slice of 0.38 μm and a 40x/1.3 oil immersion objective.

### RNA FISH (HCR *in situ* hybridization)

We used the Multiplexed HCR RNA-FISH technique ^126,127^, using reagents and adapted protocols from Molecular Instruments. The specific probe sets for each gene were designed using software from Molecular Instruments (https://www.molecularinstruments.com/). RNA FISH was then performed as follows. First, CNS from staged larvae were dissected in PBS, fixed for 20 min in 4% formaldehyde diluted in PBS, washed three times five minutes in PBS and pre-hybridised in 200 μl of hybridisation buffer for 30 min at 37°C. Meanwhile, 0.8 pmol (0.8 μl of 1 μM stock) of the probe set was added to 200 μl of probe hybridisation buffer and pre-hybridised for 30 min at 37°C. The pre-hybridisation solution was removed from the fixed samples, which were then incubated with the probe solution at 37°C overnight. Samples were washed 4 x 15 min with 500 μl warmed (37°C) wash buffer, followed by 5 min with 500 μl of 50% wash buffer/50% 5XSSC-0.1% Tween and finally by 2 x 5 min with 500 μl of 5XSSC-0.1% Tween at room temperature. Samples were further pre-amplified with 100 μl of amplification buffer for 10 min at room temperature. In the meantime, 6 pmol of hairpin h1 and 6 pmol of hairpin h2 (2 μL of 3 μM stock) were individually prepared by heating at 95°C for 90 s followed by cooling to room temperature for 30 min in the dark. Snap-cooled hairpins h1 and h2 are then added to 100 μL of amplification buffer at room temperature. The pre-amplification solution was removed from the samples, which were incubated in the hairpin solution overnight (> 12 h) in the dark at room temperature. The samples were washed 2 x 5 min, 2 x 30 min and 1 x 5 min with 5XSSC-0.1% Tween before mounting on slides. For *dpr10*, 4 pmol (4 μl of 1 μM stock) of the probe set was used instead of 0.8 pmol.

### DNA FISH

The FISH protocol was performed using oligonucleotide probes for chromosomes II and III labelled with 5′CY3 and FAM488 fluorescent dyes respectively^128^. Briefly, dissected brains were fixed for 30 min in 4% formaldehyde prepared in PBS with 0.1% tween 20, washed three times/ 10min in PBS, washed once 10min in 2xSSCT (2xSSC (Sigma S6639) + 0.1% tween-20) and once in 2xSSCT 50% formamide (Sigma 47671). For the pre-hybridisation step, CNSs were transferred to a new tube containing 2xSSCT 50% formamide pre-warmed at 92°C and denatured 3min at 92°C. For the hybridisation step, the DNA probe (40-80 ng) was prepared in hybridisation buffer (20% dextran sulphate, 2XSSCT, 50% deionised formamide (Sigma F9037), 0.5 mg.ml^−1^ salmon sperm DNA) and denatured 3 min at 92°C. Probes were added to the brains samples and hybridise 5 min at 92°C followed by overnight hybridisation at 37°C. Samples were washed with 60°C pre-warmed 2XSSCT for 10 min, washed once 5 min in 2XSSCT at RT and rinsed in PBS. CNS were mounted in Mowiol mounting medium.

### Image acquisition and processing

Confocal images were acquired using a laser scanning confocal microscope (Zeiss LSM 880, Zen software (2012 S4)) with a Plan-Apochromat 40x/1.3 Oil objective. All CNS were imaged as z-stacks with each section corresponding to 0.3-0.5 μm. Images were subsequently analysed and processed using Fiji (Schindelin, J. 2012), Volocity (6.3 Quorum technologies), and the Open-Source software Icy v2.1.4.0 (Institut Pasteur and France Bioimaging, license GPLv3). Denoising was used for some images using the Remove noise function (Fine filter) in Volocity. Images were assembled using Adobe Illustrator 28.7.1.

### Quantification of NSC number, mitotic index, mitotic phase distribution and nuclear shape

CNSs of the desired genotypes were stained with Dpn and phospho-histone H3 antibodies to detect NSC identity and mitosis, respectively. Quantification was performed on NSCs from the thoracic part of the tVNC. The Dpn signal was automatically segmented through HK-means detection on Icy and corrected manually to get NSC counts per tVNC. Mitotic phases (Prophase, Prometaphase/metaphase, Anaphase, Telophase) were manually determined by the localization and pattern of pH3 and Dpn stainings. Normalised mitotic index corresponds to the ratio between mitotic NSCs over all NSCs, then divided by the mean of this ratio for the control sample, unless stated otherwise.

The number of abnormal shapes per NSC was determined manually by the user through qualitative visual assessment. Only interphase and early prophase NSCs, identified by combined Dpn and pH3 stainings, were considered.

### Quantification of levels of RNAi knockdown by RNA FISH

For each gene, expression levels in a specific cell type were determined by calculating the total sum of RNA FISH probe intensities colocalised with the cell type compartment marker and normalising by dividing this value by the cell type’s compartment size. Each value was then normalised to the mean of the control. Colocalization was determined by the intersection of the segmented RNA FISH signal with the segmented cell type marker. Segmentation was performed in Volocity 6.3 (Quorum technologies) using adjusted protocols for intensity thresholding. Thresholds were corrected on each sample to exclude background fluorescence. The conditions for each cell type were as follows:

| Compartment | Cell type marker | Volume of calculation and reason of choice | Compartmental size and output value | Figures |
| --- | --- | --- | --- | --- |
| CG | Nrv2::GFP | - 120 x 120 x 5 $\mu\text{m}$<br>- CG layer around NSCs<br>- Manageable size for analysis | - Nrv2::GFP volume<br>- RNA FISH signal per $\mu\text{m}^3$ of CG membrane | Figures 3I, K<br>Figure S5F |
| NSC | Dpn staining | - Whole tVNC<br>- High number of NSCs | - Number of Dpn<sup>+</sup> cells<br>- RNA FISH signal per NSC | Figure 3M<br>Figure S8C |
| VNC | NA | - 120 x 120 x 5 $\mu\text{m}$<br>- Manageable size for analysis | - Volume of<br>- RNA FISH signal per $\mu\text{m}^3$ of tVNC tissue | Figure S8D |

### Quantification of spindle and DNA properties in NSCs in metaphase

CNSs of the desired conditions were stained with phospho-histone H3, α-tubulin and centrosomin (cnn) antibodies to detect metaphase DNA, central spindles and centrosomes respectively. Phalloidin was added to help identify NSC cellular borders. For all the following measures, only NSCs in late prometaphase and metaphase were considered. All analyses were performed in Icy.

Centrosome counts per NSC were determined manually using Phalloidin to circumscribe boundaries. The distance between the two centrosome poles (in μm) was calculated by tracing a line between the two cnn^+^ dots contained in one NSC.

Spindle and DNA maximal widths (in μm) were measured by drawing a line between the two furthest points on either side of the line connecting the centrosome poles. Dividing the DNA width by the spindle width gives a ratio estimating the spread of the DNA over the spindle.

DNA roundness (in %) was calculated from the threshold-based segmentation in 3D of the pH3 signal and followed the definition of the ISO 1101 standard: the normalised ratio between the radius of the minimum inscribed and largest circumscribed circles (or spheres), expressed as a percentage (100% for a circle or sphere).

Classification of spindle morphology was performed manually by scoring the number of spindles visually allocated to one of the categories by the experimenter over the total number of spindles per CNS. The classification was further refined by using 4 μm as the lower threshold for large spindle and 6 μm as the upper threshold for short spindles.

### Quantification of DNA damage by γH2Av staining

CNSs of the desired conditions were stained with Deadpan, phospho-histone H3 and γH2Av antibodies to detect NSC, mitosis and DNA damage respectively. DAPI was used to counterstain DNA. For a positive control, CNSs were incubated with aphidicolin at 100 μg.ml^-1^ for 30 min at room temperature in PBS before fixation. NSCs were segmented in 3D in Volocity 6.3 using the Dpn signal and adjusted protocols for object detection. γH2Av intensity and total DAPI intensity were determined using threshold-based segmentation in 3D, and intersected with Dpn segmentation to extract the total γH2Av intensity and of DAPI intensity for all NSCs. A value for DNA damage per NSC was then calculated as follows: ((γH2Av intensity)_TOTAL_/(DAPI intensity)_TOTAL_)/n(NSCs).

### Quantification of aneuploidy in NSC by DNA FISH

CNSs of the desired conditions were stained with probes for chromosomes II and III, and antibodies against Deadpan and phospho-histone H3. Threshold-based 3D protocols were used to segment the probes’ signals. Only NSCs in metaphase were further analysed to ensure all NSCs had passed replication, and the expected ploidy was comparable across conditions and were identified by the combination of Dpn and pH3 stainings. The sum of intensities of all segmented objects within one Dpn^+^-segmented cell was determined for each probe as an indicator of DNA content for this locus per NSC.

### Measure of membrane proximity by split-GFP

To determine the NSC-CG membranes proximity, the nuclei area was segmented using surface segmentation from Imaris on the Dpn signal. The resulting nuclei mask M_nuclei_ was first dilated with an element of size 11×11×3 pixels generating the mask M_dilate_, in order to extend the considered area to include the area corresponding to the NSCs membrane. M_nuclei_ was also eroded with an element of size 3×3×1 pixels to generate the mask M_erode_. The membrane mask M_membrane_ was computed using the difference between M_dilate_ and M_erode_. Any region not containing the *LexAop-myr-tomato* signal was removed from the quantification. We then computed an intensity ratio of the split-GFP signal by comparing the number of pixel higher than an Otsu threshold M_dilate_ ROI, with the number of pixels contained in the M_membrane_ ROI. This ratio is called GFP intensity average and is expressed in Arbitrary Unit (AU). The Python quantification script was based on the scikit-image library^129^.

### Measure of NSC membrane geometry

To determine the geometric parameters of the NSC membrane between control and *dpr10* RNAi conditions from the SPLIT experiment, we manually drew the contours of the *grh > myr-tdTomato* signal around one NSC (marked by Dpn) using the polygon tool from Icy. An average of 20 NSCs in interphase or early prophase was sampled per tVNC. The convexity of each individual contour was then determined using the ROI statistics function of Icy, which defines convexity as the ratio between the volume and the volume of the smallest convex envelope containing the ROI, expressed as a percentage (a convex object has a value of 100 %).

### Quantification of nuclear lamina deformation and lamin levels in NSC

CNSs of the desired conditions were stained with Dpn, phospho-histone H3 and Lamin Dm0 (Lam) antibodies to detect NSC identity, mitosis and nuclear lamin/B-type lamin respectively. For each CNS, an average of 35 NSCs in interphase or early prophase was cropped at the equatorial plane. On each cropped image, a median filter of size 4 and a Gaussian filter (sigma = 1) were applied before segmentation using the Python version of cellpose ^130^, each on both Lam and Dpn channels. Segmented objects in the border of the image were discarded and only the largest object was selected for quantification. The inner boundary of Lam was detected using Otsu thresholding applied on the whole crop. We computed 4 markers in the quantification step: the proportion of Lam signal reported to the total area of the nuclei and its total fluorescence normalised by the size of the nuclei; the proportion of Dpn signal reported to the total area of the nuclei (based on Lam segmentation); and using both Lam and Dpn signals, we quantified the size of the overlap between the two signals reported to the total area and then proportion of pixels not segmented with Dpn nor inner boundary of Lam and refered hereafter as "holes". Some images were discarded from the analysis using the following thresholds: proportion of Dpn signal related to outer Lam higher than 1, overlap between Deadpan and outer Lam lower than 0.1 or proportion of holes higher than 0.25. Statistical comparisons were then performed using linear mixed models with R software (lme4^131^ and emmeans^132^).

### Live imaging of neural stem cell divisions and CRISPR knock-in lines

CNS were dissected in freshly prepared live imaging medium composed of Schneider’s medium (ThermoFisher 21720024), 5% Fetal Bovine Serum (Sigma F4135), 1/100 larval lysate and 1/100 Penicillin/Streptomycin (ThermoFisher 15140). The larval lysate was prepared with 20 larvae mixed in 200 μl of Schneider’s until obtaining a whole homogenate that was spun down at 6000 rpm for 5 minutes at 4°C and the supernatant was recovered. 15 mm glass bottom dishes were previously coated with 1% low melting point agarose (ThermoFisher 16520100) in Schneider’s with 1/100 Penicillin/Streptomycin. Wells of approximately 2 mm diameter were made in the solidified agarose and the surface was filled with live imaging medium. Dissected CNS were loaded in the wells and oriented with the ventral side facing down, a lid was placed on the top of the dish. After a couple of minutes, the CNS were settled down and the sample ready to image. Live imaging was performed with a temperature-controlled environmental chamber set at 25°C and a 40x/1.3 oil immersion objective on a LSM880 confocal microscope. Images were collected at 1024×1024 pixels following time series of 60 s interval.

### Measure and statistics of cell cycle length in neural stem cells

Live imaging data was manually analysed to count the length of NSCs mitosis (considering time 0 the first observation of clear chromatin condensation with a *H2Av∷mRFP* reporter).

To compare the mitotic length of our control and *dpr10* knockdown conditions, we first plotted on a histogram the distribution of the NSCs division times in both groups. Overall, distributions in both groups looked highly similar except for a few outliers. Then, we tested for the existence of outliers in both groups by assuming that variation in mitotic length could be suitably captured by an exponentially modified Gaussian distribution (henceforth, exGaussian distribution). This distribution was considered flexible enough to model intermitotic times in a wide range of cells and conditions^133^ with only three parameters (namely, the mean, the standard deviation as well as the exponential rate).

We sampled a particular value for each of the parameters where the probability of sampling a particular combination of three was proportional to the likelihood of the corresponding distribution fitting the observed values of the control group (Metropolis-Hastings algorithm). Based on the sampled parameter values, the number of values corresponding to the size of each group (98 for control and 162 for *dpr10* knockdown) was randomly obtained from the theoretical exGaussian distribution. The highest value was recorded in each set of simulated values (control and *dpr10* knockdown group). This step was repeated 100 times (and in each group) so that 100 "highest values" were recorded for a particular set of parameter values (mean, standard deviation, rate). After repeating the procedure 20 000 times, p-values were estimated for each observed value as: the number of times the simulated "highest values" exceeded the given observed value, divided by the total number of repetitions (*i.e.*, 100 x 20,000). Each single value could be considered a significant outlier whenever a p-value < 0.05.

### Photo-ablation ablation and measure of cortical tension in cortex glia membrane

CNS were prepared for live-imaging as detailed above. Photo-ablation coupled with live-imaging with a 40X oil objective (Nikon plan fluor, N.A. 1.30) was performed as described in ^134^ using a pulsed UV-laser (355nm, Teem photonics, 20kHz, peak power 0.7kW) coupled to a Ilas-pulse module (Gataca systems) attached to a spinning disk microscope (Inverted Nikon Eclipse). Nrv2::GFP was used to visualize the cell cortex of the CG. The CG network is dynamic and likely heterogeneous in tension. To minimize variance in laser ablation, we restricted the cutting area to the most anterior part of the tVNC and targeted only membranes clearly in contact with NSCs in close range of their equatorial plane. Photo-ablation was achieved at 355 nm using 55% power to minimize cavitation, and with a cut thickness of 1 μm. Images were taken every 200 ms (frequency of 5 Hz) and photo-ablation happened 10 images/2 s after the beginning of the movie with 40 repetitions over an exposure time of 100 ms. Laser for imaging was the same used at 35 % of power. Imaging was performed at 488nm at a power minimizing photobleaching.

The resulting movies were curated to remove cavitation events and clear failure in ablation. One recognizable landmark in the Nrv2::GFP signal (vertices; bulges) was then selected on each side of the laser cut and outside of the burn zone for each remaining movie. These two landmarks (1 ; 2) were tracked over time and their spatial x,y coordinates recorded. The Euclidian distance between these two landmarks was calculated using the following formula: ((x2-x1)^2^+ (y2-y1)^2^)^0.5^ and plotted over time. All values were offset by the mean of the first nine timepoints (before ablation).

Several curve fittings were then tested, based on the core formula from ^135^, and after multiple tests, a combined linear + exponential model was deemed the best fit, with the following formula:

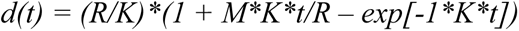

where

d(t) corresponds to the strain.

R is related to the tensile force present at the cut location before ablation.s K is linked to the elasticity.

M is the linear term determined by curve fitting.

The gradient or initial recoil is d(y)/d(t) at t = 0 (d(t_0_)) and equal to M + R.

The M, R, K and d(t_0_) values were then calculated for each movie (n(control) = 18; n(ROKCA) = 18; and n(*dpr6*+*dpr10* RNAis) = 23). Outliers were identified with the ROUT method (Q = 1%) and a Kruskal-Wallis test was performed on the R values of the remaining movies (n(control) = 15; n(ROKCA) = 15; and n(*dpr6*+*dpr10* RNAis) = 29) to determine the significance.

### Statistical tests and experimental reproducibility

Statistical tests used for each experiment are stated in the figure legends. All statistical tests were performed using GraphPad Prism 10.3.1 or in R for the Linear Mixed Model.

Mitotic index, RNA FISH signals and γH2Av expression were normalised to the mean of the control condition for each replicate in order to account for replication effect. For most experiments, at least two independent experimental replicates, each providing at least six biological replicates (tVNC, with an average of 155 NSCs each), were performed and analyzed for reproducibility through a Nested analysis on Prism (Nested t-test if two conditions and Nested one-way ANOVA if more than two conditions). If the replicates were detected as statistically different, then the statistical significance of the whole dataset is given by the p-values of the Nested analysis or assessed through a linear mixed model. If the replicates were not deemed statistically different, then grouped analyses were then performed. For comparing two conditions, an unpaired Student t-test was used when they passed the normality test (D’Agostino and Pearson test) and Mann-Whitney U test when they did not pass the normality test. For comparing more than two conditions, a one-way ANOVA with Tukey’s multiple comparisons test was used on normal data and a Kruskal–Wallis H test with Dunn’s multiple comparisons test for data which did not pass normality.

For lamin surface area and folding, spindle features and DNA roundness, each CNS was sampled by at least 30 crops and was considered as one biological replicate. Regardless of the chosen representation (box or scatter dot plots), the analysis accounted for the sampling size of each CNS while comparing differences between them, using either a nested analysis or a linear mixed model.

For DNA FISH analysis, between 10 and 15 NSCs in metaphase were analysed per CNS, with a minimum of 8 CNSs per condition. The analysis accounted for the sampling size of each CNS while comparing differences between them, using either a nested analysis or a linear mixed model.

For all knockdown (RNAi) and misexpression experiments, penetrance and expressivity are represented through sample distribution for the parameter assessed, with individual values (dots) placing each biological replicate with respect to control’s distribution and mean. n numbers are indicated in the figure legends.

### Data representation

All images displayed are representative.

Box plots show minimal value (bottom whisker), first quartile (25th percentile, lower limit of the box), a median of the interquartile range (middle horizontal line), third quartile (75th percentile, the upper limit of the box) and maximal value (top whisker). Individual values are superimposed and colour-coded to represent multiple experimental replicates. In addition, shading in the grey range indicate replicates not statistically different, while khaki colouring is used to highlight statistically different groups.

Stacked bars were generated using the mean for each phase distribution. Error bars represent the Standard Error of the Mean (SEM).

For dot plots, the identity of the dots is described in the legends of the figure and the horizontal line corresponds to the median.

Significant and of interest p-values are displayed directly on the graph, together with their usual abbreviations (ns, p≥0.05; *, p < 0.05; **, p <0.01; ***, p<0.001; ****, p<0.0001) to simplify interpretation.

## Acknowledgments

We thank members of the Spéder lab for helpful discussion and suggestions; P. Campagne for the analysis of the NSC mitotic length; A. Joudat for sharing her expertise on the generation of the fluorescently tagged CRISPR/Cas9-engineered lines; R. Levayer for input on the modulation of cortical tension; F. Murphy for helping with and writing the code for recoil curve fitting. We thank L. Zipursky, C. González, K. Scott, the VDRC, BDSC and DSHB stock centers for reagents. This work has been funded by a starting package from Institut Pasteur/LabEx Revive (ANR-10-LABX-0073) to P.S., a JCJC grant from Agence Nationale de la Recherche (NeuraSteNic, ANR-17-CE13-0010-01) to P.S, a Tremplin ERC grant from Agence Nationale de la Recherche (SiStemNiC, ANR-22-ERCC-0005) and a Projet Fondation ARC from the Association pour la Recherche contre le Cancer to P.S. We acknowledge the support of France-BioImaging (ANR-10-INBS-04) for this work.

## Author Contributions

A.S-C and P.S. conceived the experimental design. A.S-C, P.S. and D.B. analyzed most data. A.S-C. performed experiments of Figures 1; S1; 2; S2; 3A-C, N-P; S3B,D,E-G; 4; S4A-E; 5D-G and H-K; S5A,F-H; 6E-I; S6A-C; S7H-K and S8A. D.B. performed experiments of Figures 3H-M; S3A, C, H; S4I-K; 5D-G; S5B-E, H and S8B-D. P.S. performed experiments of Figures S4F-H; 5D-G; 6A-I; S6A-C; S7A-G; 7 and S8E-L and analysed the raw TaDa-PolII data. A.K. performed the laser ablation experiments (Figures 5A-C). S. R. performed the Split-GFP analysis (Figure 3B-C). E.P. performed the analysis of Lamin area and levels (Figures 7B-C, E-F). J-Y.T. provided supervision for the image analysis components of the study. A.B-L helped with the TaDa-PolII experiment (Figure 1). N.D.L.O. performed the clustering on the TaDa-PolII datasets. L.V. helped setting up the experimental conditions for and advised on the laser ablation experiments (Figures 5A-C). Y.B. provided expertise on tension manipulation and the generation of the CRISPR/Cas9-based lines as well as unpublished transgenic lines (Figures 3N-P and S3F-H). A.S-C. and P.S. wrote the manuscript.

## Data availability statement

The TaDa-PolII dataset is available as Table S1. The scripts for Lamin analysis (10.5281/zenodo.14358623) and recoil curve fitting (10.5281/zenodo.14344916) will be accessible on Zenodo. Quantifications are available in the Data source spreadsheet. The image datasets generated during and/or analysed during the current study are available from the corresponding author on reasonable request.

---

**Table S1. TaDa-Poll gene clusters**

(Clusters tab). The Cluster column corresponds to the identity of the cluster to which the gene belongs. MeanT1, MeanT2 and MeanT3 correspond to the mean log2 ratio change for binding across the annotated gene (DamPol II/Dam) and biological replicates at T1, T2 and T3 timepoints respectively.

(FDR values tab). The nomenclature Tx_y is used to represent the different biological replicates, with: x = timepoint and y = replicate. The value of the FDR (False Discovery Rate) is written for each gene for each biological replicate. Its value will determine whether the gene is considered as expressed (“1”, FDR < 0.01) or not (“0”, FDR ≥ 0.01) at each timepoint.

**Table S2. Drosophila genotypes, crosses and culture regimens per experiment**

**Movie S1. Neural stem cell division in control conditions**

Confocal time-lapse video of a control tVNCs (*cyp > -*). Ubiquitous expression of *H2Av∷mRFP* marks chromatin in the tissue (magenta) and *nrv2::GFP* (green) labels CG membranes. Larvae are observed at ALH60 at 29°C. The NSC goes periodically out of the plane of the movie due to some contractile movements of the tissue in the live-imaging set-up.

**Movie S2. Neural stem cell division under *dpr10* knockdown in the cortex glia**

Confocal time-lapse video of one NSC under *dpr10* knockdown in CG (*dpr10 RNAi^VDRC103511^*). Ubiquitous expression of *H2Av∷mRFP* marks condensed chromatin in the tissue (magenta) and *nrv2::GFP* (green) labels CG membranes. Larvae are observed at ALH60 at 29°C.

**Movie S3. Neural stem cell divisions under *dpr10* knockdown in the cortex glia**

Confocal time-lapse video of a tVNC under *dpr10* knockdown in CG (*dpr10 RNAi^VDRC103511^*). Ubiquitous expression of *H2Av∷mRFP* marks condensed chromatin in the tissue (magenta) and *nrv2::GFP* (green) labels CG membranes. Larvae are observed at ALH60 at 29°C. Several NSC divisions can be observed in the movie. Cyan circles show examples of NSCs undergoing mitosis in a normal timeframe and white circles show examples of NSCs being arrested for several hours in metaphase.

**Movie S4. Dynamic punctuated expression of mKate2**∷**Dpr10 along CG membrane**s

Confocal time-lapse video of a tVNC of a CRISPR/Cas9 engineered line *mKate2::dpr10* (magenta) and *nrv2::GFP* (green). *mKate2::dpr10* shows a punctuated expression that colocalizes partly with CG membrane. Larvae are observed at ALH72 at 25°C.

**Movie S5. Colocalization between mKate2∷Dpr10 and Dpr10::GFP**

Confocal time-lapse video of a tVNC of a CRISPR/Cas9 engineered line *mKate2::dpr10* (magenta) and *dpr10::GFP* (green). *mKate2::dpr10* and *dpr10::GFP* colocalize on the dotted pattern. Larvae are observed at ALH72 at 25°C.

**Movie S6. Punctuated expression of DIP-α∷GFP along NSC membranes**

Confocal time-lapse videos of a tVNC of a CRISPR/Cas9 engineered line *DIP-α::GFP* (green) and grh-*myr-mScarlet* (magenta). *DIP-α::GFP* is expressed in neurons and colocalizes also with NSC membrane marker. Larvae are observed at ALH72 at 25°C.

**Movie S7. Co-expression of mKate2::Dpr10 and DIP-α∷GFP**

Confocal z-stack videos acquired live of a tVNC where both *mKate2::dpr10* (magenta) and *DIP-α::GFP* (green) are expressed. Some dynamic co-localization can be seen between the two markers. Several mKate2::Dpr10 dots are visible without DIP-α::GFP colocalization (magenta arrowheads), before becoming charged and colocalizing with DIP-α::GFP (white arrowheads). The magenta circle shows a mKate2::Dpr10 dot which does not colocalize with DIP-α::GFP during the timeframe of the movie. Larvae are observed at ALH72 at 25°C.

**Movie S8. Recoil velocity in cortex glia membrane following laser ablation in control condition.**

Representative movie (average recoil) for *cyp > -*. CG membrane is marked with Nrv2::GFP (grey). The cyan arrowhead marks the place and time of laser ablation. Larvae are observed at ALH66 at 29°C.

**Movie S9. Recoil velocity in cortex glia membrane following laser ablation under constitutive activation of actomyosin contractility.**

Representative movie (average recoil) for *cyp > ROK^CA^*. CG membrane is marked with Nrv2::GFP (grey). The cyan arrowhead marks the place and time of laser ablation. Larvae are observed at ALH66 at 29°C.

**Movie S10. Recoil velocity in cortex glia membrane following laser ablation under double dpr10 and dpr6 knockdowns.**

Representative movie (average recoil) for *cyp > dpr6 RNAi^VDRC41161^ + dpr10 RNAi^BDSC27991^*. CG membrane is marked with Nrv2::GFP (grey). The cyan arrowhead marks the place and time of laser ablation. Larvae are observed at ALH66 at 29°C.

