## Supplemental Figures and Legends for "Adhesion-Controlled Mechanics of the Glial Niche Regulate Neural Stem Cell Proliferative Potential"

Segú Cristina et al, Figure S1

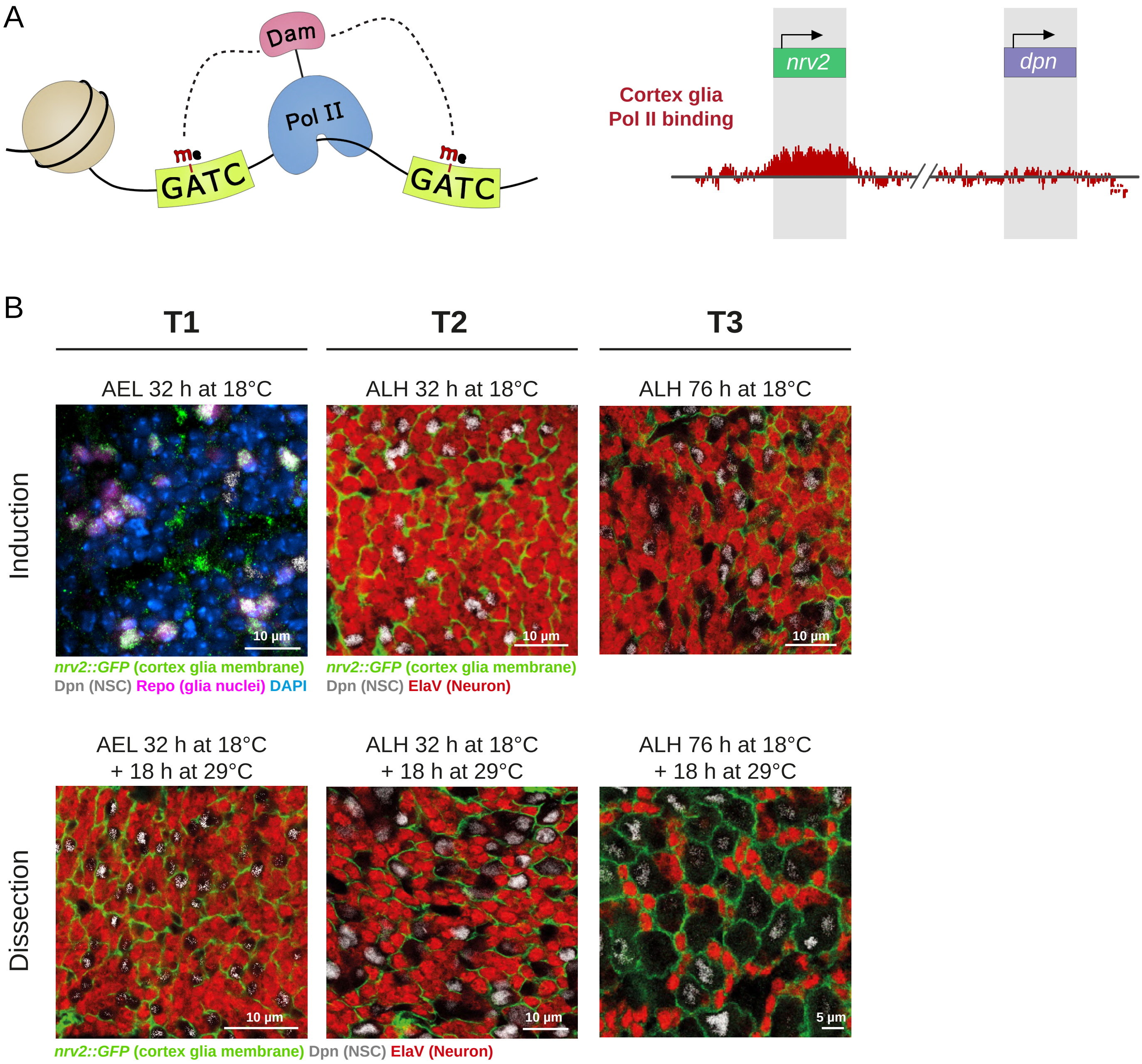

**Figure S1. Transcriptional analysis of the cortex glia using the Targeted DamID (TaDa)-PolII technique, related to Figure 1**

(A) Schematic of the principle of methylation-based recording of Polymerase II binding. The fused Dam methylates GATC sequences around the sites where Pol II binds to the DNA at the start and along transcription. CG transcriptional profile can be extracted from the methylation profile on the genome. Transcribed genes, such as *nrv2* (expressed in CG), show high Pol II occupancy, while non-transcribed genes, such as *dpn* (specific to NSCs), show low occupancy. (B) Confocal images of the timepoints chosen to start and stop the induction of the TaDa PolII to record CG gene expression at T1 (before nutrition-induced membrane growth), T2 (during membrane growth) and T3 (at the time of NSC encasing). AEL: after egg laying. ALH: after larval hatching. CG membrane: Nrv2::GFP (green), NSC: Dpn (grey), neuron: ElaV (red), glia: Repo (magenta). DAPI is used to counterstain the nuclei.

See also Tables S1 and S2.

Segú Cristina et al, Figure S2

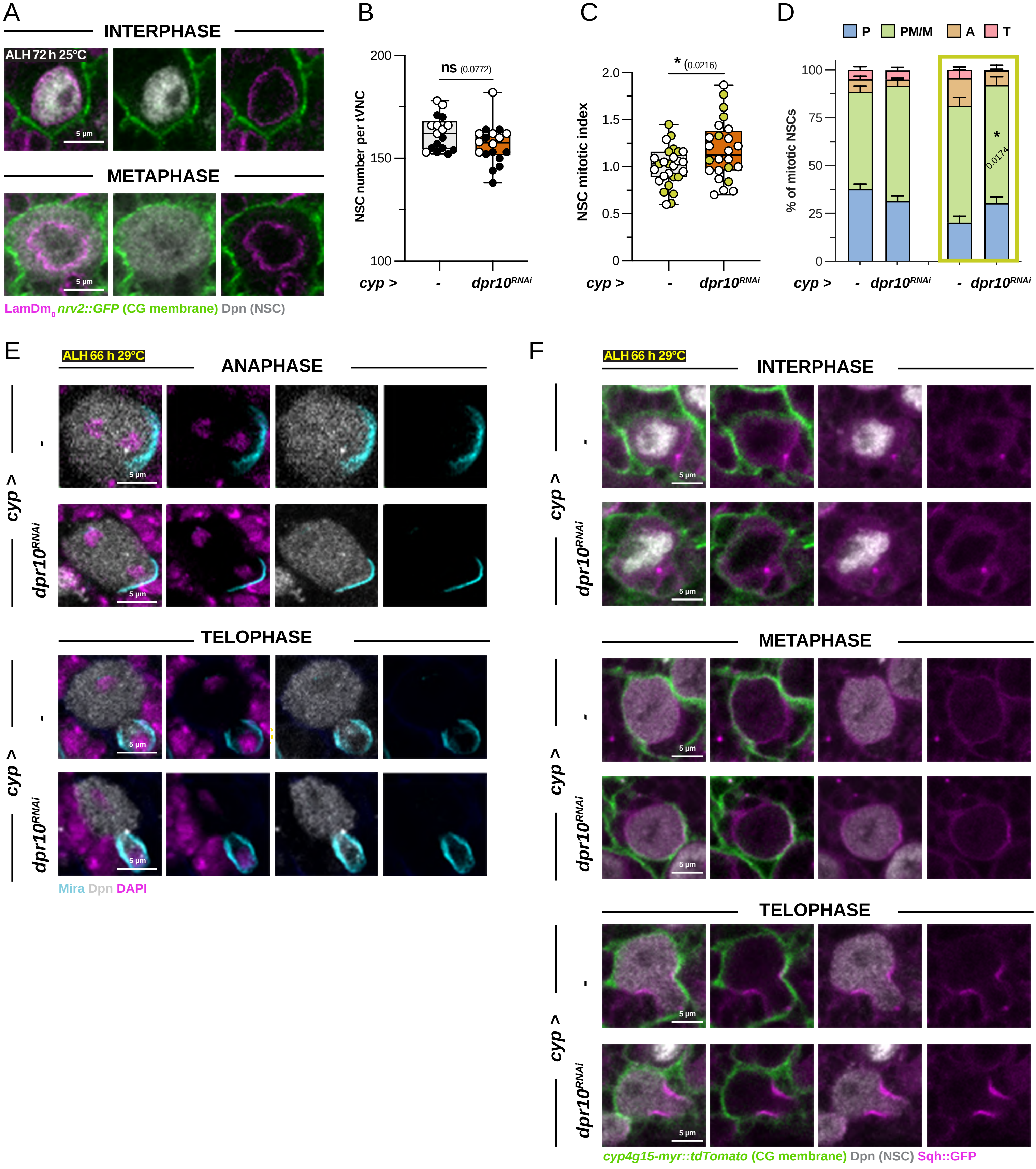

**Figure S2. *dpr10* is required non-autonomously in the cortex glia to control neural stem cell division but is dispensable for asymmetric polarity, related to Figure 2**

(A) Confocal images of NSCs stained with LamDm0 (magenta) to mark nuclear lamina, and with Dpn (grey) for NSC nuclei. CG membranes express Nrv2::GFP (green). Dpn antibody can be used as a proxy of nuclear shape during interphase. Dpn, a transcription factor in NSCs, is confined within the nuclear envelope during interphase and prophase. In prometaphase, the nuclear envelope becomes more permeable during the semi-closed mitotic process taking place in *Drosophila* NSCs and Dpn disperses and occupies the entire cytoplasm.

(B) Box plot of NSC numbers in *cyp* > - (n = 17 tVNCs) and *cyp* > *dpr10* $RNAi^{VDRC103511}$ (n = 18 tVNCs). Unpaired Student t-test.

(C-D) Box plot of mitotic index (normalized to control) and stacked bar chart of phase distribution in NSCs for *cyp* > - and *cyp* > *dpr10* $RNAi^{VDRC103511}$. Two replicates are shown with either white (n = 17 tVNCs, and control (n = 21 tVNCs)) or khaki (n = 7 tVNCs, and control (n = 6 tVNCs)) dots and frames. Both experimental replicates show a higher mitotic index, while only the khaki-coloured replicate shows a higher percentage of NSCs in prometaphase/metaphase. Unpaired Student t-test (C) and two-way ANOVA with Šídák multiple comparison test (D).

(E) Confocal images of NSCs stained for Miranda (Mira, cyan) to follow asymmetric division. Asymmetric localization of Miranda happens similarly in the control (*cyp* > -) and under *dpr10* knockdown in the CG (*cyp* > *dpr10* $RNAi^{VDRC103511}$), at both anaphase and telophase.

(F) Confocal images of NSCs expressing a ubiquitous *sqh::GFP* reporter of myosin II (magenta). Myosin II was similarly expressed along the cell cycle in control (*cyp* > -) and under *dpr10* knockdown in the CG (*cyp* > *dpr10* $RNAi^{VDRC103511}$), including the characteristic localization at the cytokinesis site during telophase.

For box plots, individual values are superimposed and colour-coded to represent multiple experimental replicates.

ns, $p \geq 0.05$; *, $p < 0.05$; **, $p < 0.01$; ***, $p < 0.001$; ****, $p < 0.0001$

P: Prophase, PM/M: Prometaphase/metaphase, A: Anaphase, T: Telophase.

See also Table S2.

Segú Cristina et al, Figure S3

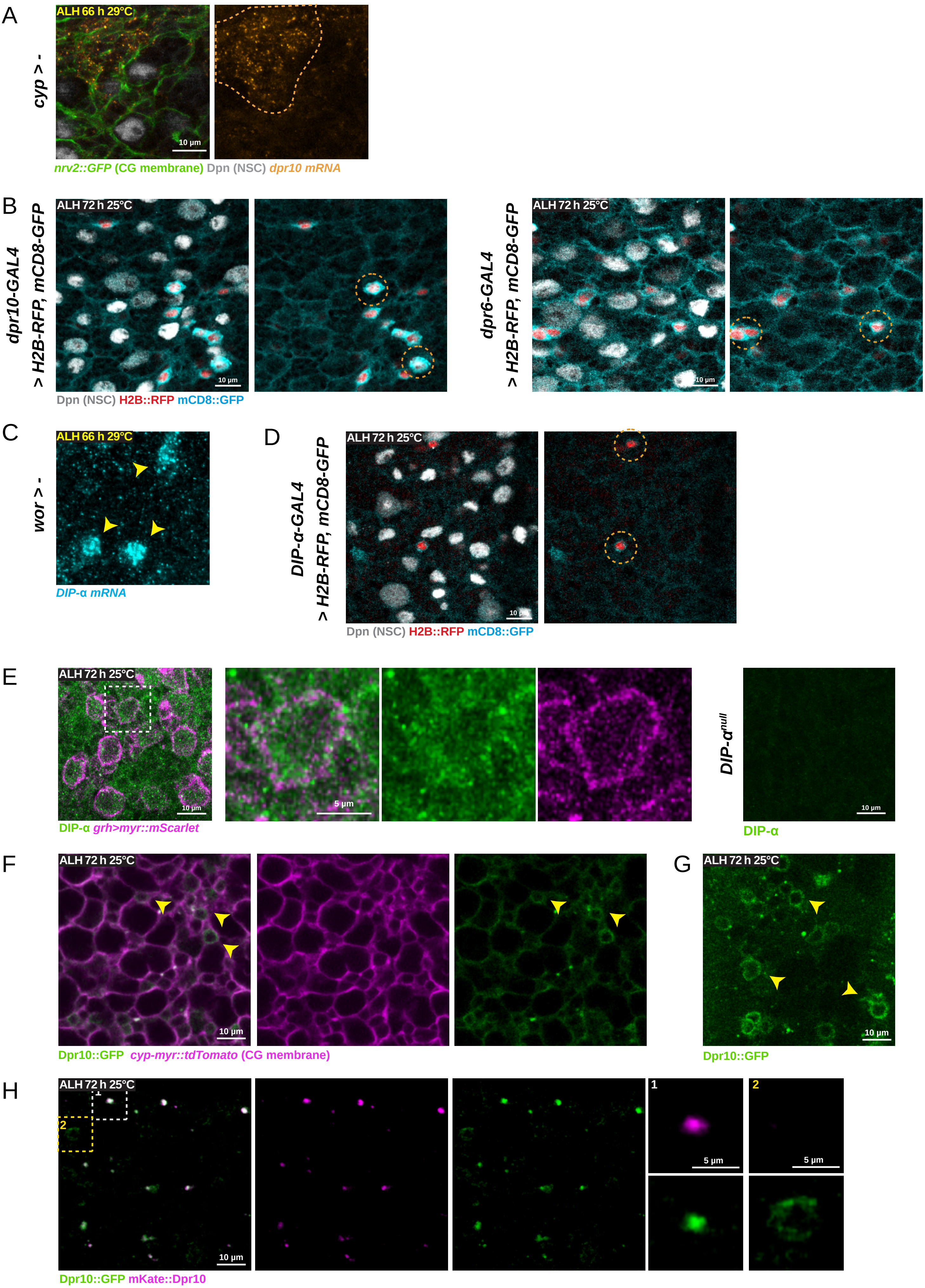

**Figure S3. Dpr10 forms an Ig-based interaction triad with DIP-α and Dpr6 in the neurogenic niche, related to Figure 3**

(A) Confocal image of a close-up of a control tVNC showing *dpr10* RNA expression in neurons (dashed orange area).

(B) Confocal images of a tVNC of *dpr10-GAL4* and *dpr6-GAL4* driving *H2B::RFP* (red) and *mCD8-GFP* (cyan) and stained with Dpn (grey) to mark the NSC nuclei. Both *dpr10-GAL4* and *dpr6-GAL4* show neuronal expression and a pattern reminiscent of CG structure, with a grid-type membrane pattern. Dashed orange circles show examples of stained neurons.

(C) Confocal images of a close-up of a control tVNC showing *DIP-α* mRNA signal in neurons (yellow arrowheads).

(D) Confocal images of a tVNC of *DIP-α-GAL4* driving *H2B::RFP* (red) and *mCD8-GFP* (cyan) and stained with Dpn (grey) to mark the NSC nuclei. *DIP-α-GAL4* shows some neuronal expression but does not appear to drive in NSCs. Dashed orange circles show examples of stained neurons.

(E) Confocal images of a tVNC stained with a DIP-α antibody (green) and either expressing a marker for NSC membranes (*grh*$^{D4}$*-myr::mScarlet*, magenta) or in a *DIP-α*$^{null2}$ mutant conditions. DIP-α is expressed within the NSC niche, including in NSCs and its expression disappears in *DIP-α*$^{null2}$ mutants. Close-ups of a NSC highlights the expression of DIP-α over the NSC membrane.

(F) Confocal image of a CRISPR/Cas9 engineered line *dpr10::GFP* (green) crossed to a line where the CG membrane is marked (*cyp4g15-myr::mtdTomato*, magenta). Yellow arrowheads indicate neuronal expression.

(G) Confocal image of a CRISPR/Cas9 engineered line *dpr10::GFP*, below the NSC plane. Yellow arrowheads indicate neuronal expression.

(H) Still image of a time-lapse movie (Movie S5) of the CRISPR/Cas9 *mKate2::dpr10* (magenta) and *dpr10::GFP* (green) together. Insert 1 (white frame) highlights the co-localization between these two fusion proteins on the dotted pattern. Insert 2 (yellow frame) highlights the exclusive localization of a continuous membrane *dpr10::GFP* signal in neuron.

See also Table S2.

Segú Cristina et al, Figure S4

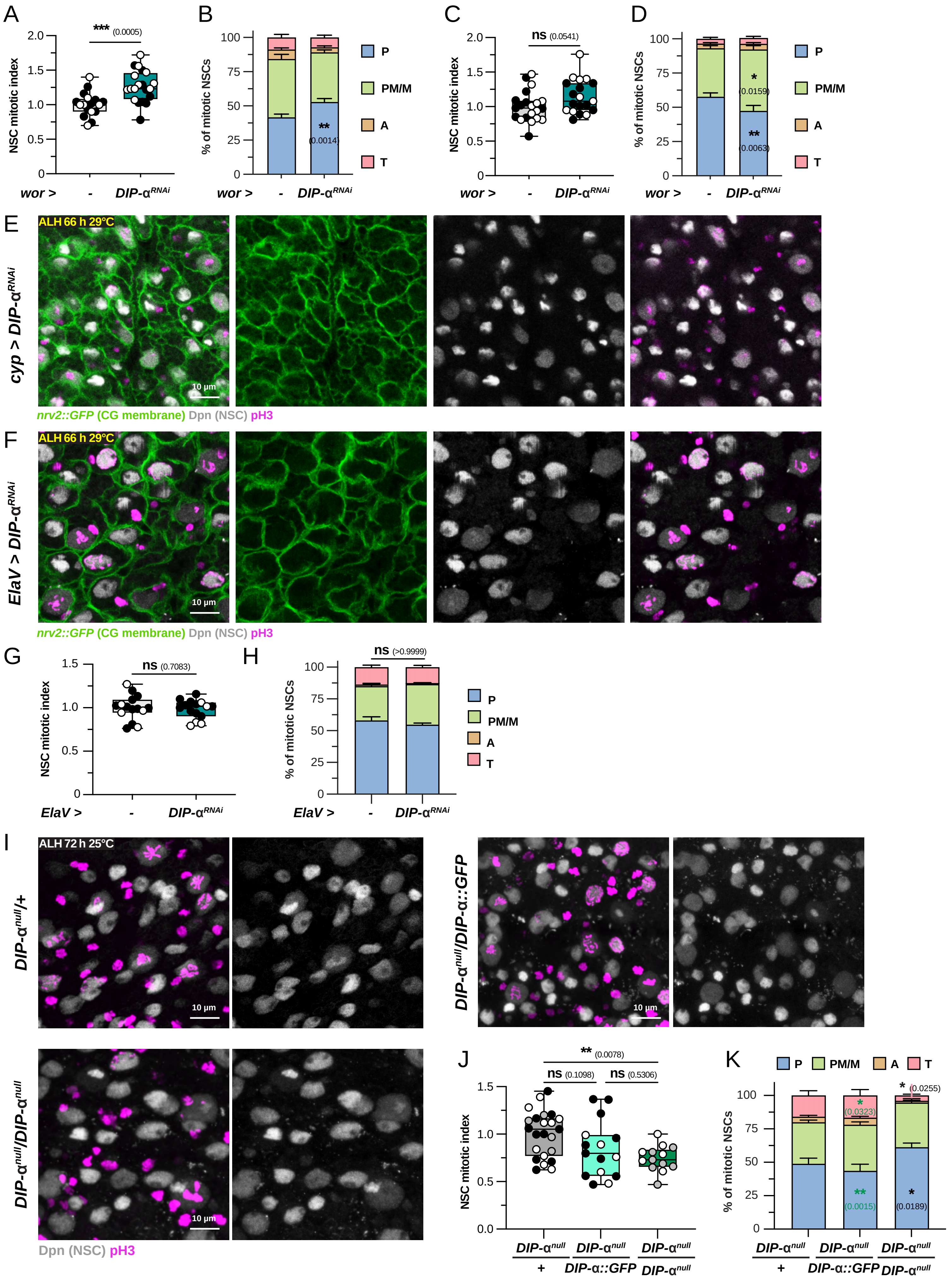

**Figure S4. DIP-α is the partner of Dpr10 in the neurogenic niche, related to Figure 4**

(A-B) Box plot of mitotic index (normalized to control) and stacked bar chart of phase distribution in NSCs for *wor* > - (n = 17 tVNCs) and *wor* > *DIP-α RNAi*$^{VDRC104044}$ (n = 20 tVNCs). This experimental replicate shows a higher mitotic index coupled with higher percentage of NSCs in prophase. Unpaired Student t-test (A) and two-way ANOVA with Šídák multiple comparison test (B).

(C-D) Box plot of mitotic index (normalized to control) and stacked bar chart of phase distribution in NSCs for *wor* > - (n = 21 tVNCs) and *wor* > *DIP-α RNAi*$^{VDRC104044}$ (n = 19 tVNCs). This experimental replicate shows a higher mitotic index coupled with higher percentage of NSCs in prometaphase/metaphase. Unpaired Student t-test (C) and two-way ANOVA with Šídák multiple comparison test (D).

(E) Confocal images of the tVNC under *DIP-α* knockdown in the CG (*cyp* > *DIP-α RNAi*$^{VDRC104044}$*)* showing CG membranes and NSC proliferation.

(F) Confocal images of the tVNC under *DIP-α* knockdown in neurons (*ElaV* > *DIP-α RNAi*$^{VDRC104044}$) showing CG membranes and NSC proliferation.

(G-H) Box plot of mitotic index (normalized to control) and stacked bar chart of phase distribution in NSCs for *ElaV* > - (n = 15 tVNCs) and *ElaV* > *DIP-α RNAi*$^{VDRC104044}$ (n = 15 tVNCs). Unpaired Student t-test (G) and two-way ANOVA with Šídák multiple comparison test (H).

(I) Confocal images of the tVNCs of heterozygous *DIP-α*$^{null2}$ mutants (*DIP-α*$^{null2}$/+), homozygous *DIP-α*$^{null2}$ mutants (*DIP-α*$^{null2}$/*DIP-α*$^{null2}$) and *DIP-α*$^{null2}$ mutant over the *DIP-α::GFP* CRISPR line (*DIP-α*$^{null2}$/*DIP-α::GFP*) showing NSC proliferation.

(J-K) Box plot of mitotic index (normalized to heterozygous *DIP-α*$^{null2}$ mutants) and stacked bar chart of phase distribution in NSCs for *DIP-α*$^{null2}$/+ (n = 23 tVNCs), *DIP-α*$^{null2}$/*DIP-α*$^{null2}$ (n = 13 tVNCs) and *DIP-α*$^{null2}$/*DIP-α::GFP* (n = 15 tVNCs). One-way ANOVA (J) and two-way ANOVA with Šídák multiple comparison test (K).

For box plots, individual values are superimposed and colour-coded to represent multiple experimental replicates.

ns, $p \geq 0.05$; *, $p < 0.05$; **, $p < 0.01$; ***, $p < 0.001$; ****, $p < 0.0001$

P: Prophase, PM/M: Prometaphase/metaphase, A: Anaphase, T: Telophase.

See also Table S2.

Segú Cristina et al, Figure S5

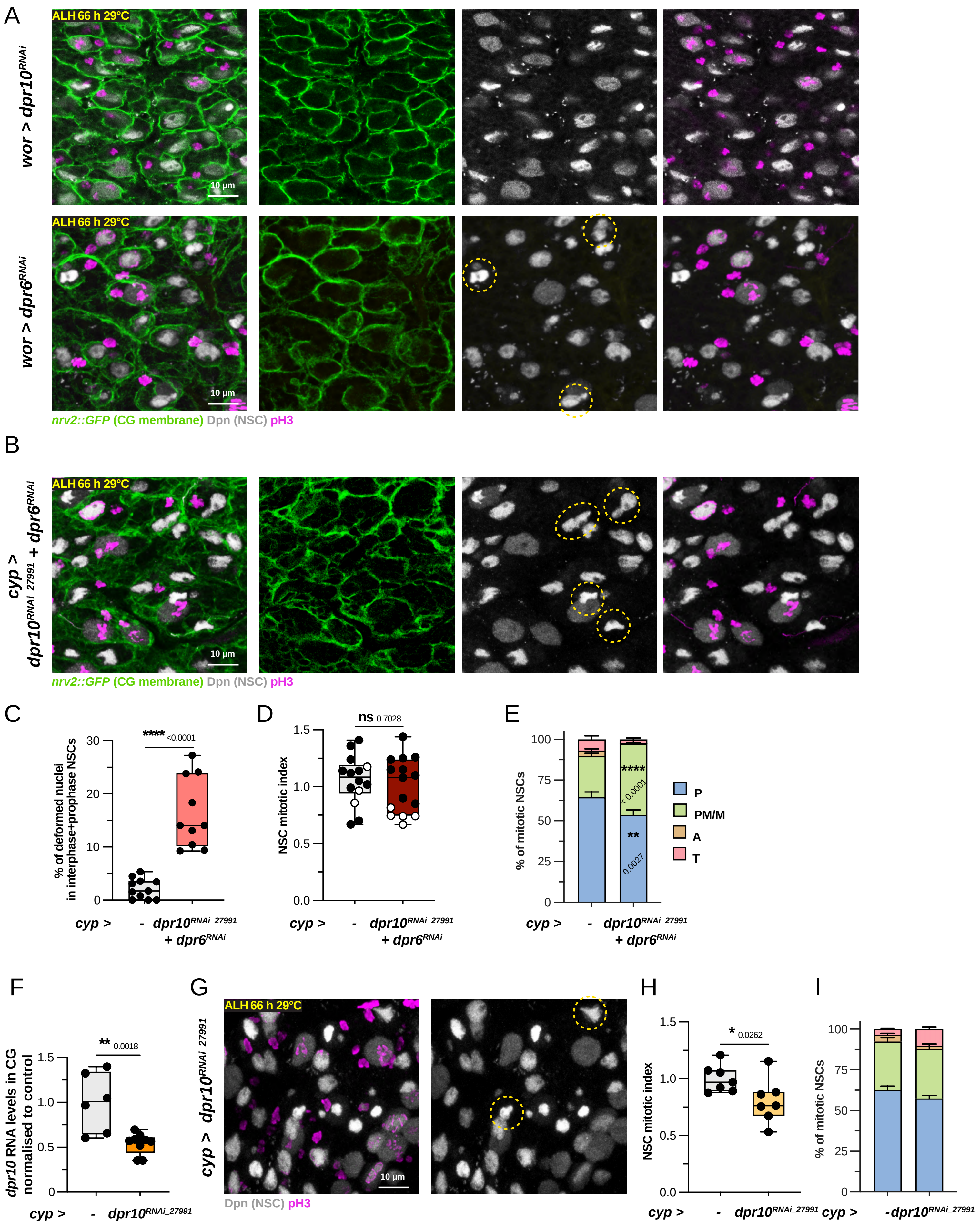

**Figure S5. Dpr10 forms an Ig-based interaction triad with DIP-α and Dpr6 in the neurogenic niche, related to Figure 4**

(A) Confocal images of a tVNC under either *dpr10* (*wor > dpr10 RNAi*$^{VDRC103511}$) or *dpr6* (*wor > dpr6 RNAi*$^{VDRC41161}$) knockdown in NSCs (*wor > dpr6 RNAi*$^{VDRC41161}$).

(B) Confocal images of a tVNC under double *dpr10* and *dpr6* knockdowns in CG (*cyp > dpr6 RNAi*$^{VDRC41161}$ + *dpr10 RNAi*$^{BDSC27991}$).

(C) Box plot of the percentage of NSCs (in interphase + early prophase) showing deformed nuclear shapes in *cyp* > - (n = 11 tVNCs) and *cyp > dpr6 RNAi*$^{VDRC41161}$ + *dpr10 RNAi*$^{BDSC27991}$ (n = 10 tVNCs). Unpaired Student t-test.

(D-E) Box plot of mitotic index (normalized to control) and stacked bar chart of phase distribution in NSCs for *cyp* > - (n = 14 tVNCs) and *cyp > dpr6 RNAi*$^{VDRC41161}$ + *dpr10 RNAi*$^{BDSC27991}$ (n = 15 tVNCs). Nested t-test (D) and two-way ANOVA with Šídák multiple comparison test (E).

(F) Box plot of *dpr10* RNA levels in the CG for *cyp*>- (n = 6 tVNCs) and *cyp> dpr10 RNAi*$^{BDSC27991}$ (n = 9 tVNCs). Unpaired Student's t-test.

(G) Confocal images of a tVNC under *dpr10* knockdown in CG with the RNAi line BDSC 27991 (*cyp > dpr10 RNAi*$^{BDSC27991}$).

(H-I) Box plot of mitotic index (normalized to control) and stacked bar chart of phase distribution in NSCs for *cyp* > - (n = 7 tVNCs) and *cyp > dpr10 RNAi*$^{BDSC27991}$ (n = 7 tVNCs). Mann-Whitney U test (H) and two-way ANOVA with Šídák multiple comparison test (I).

For box plots, individual values are superimposed and colour-coded to represent multiple experimental replicates.

ns, $p \geq 0.05$; *, $p < 0.05$; **, $p < 0.01$; ***, $p < 0.001$; ****, $p < 0.0001$

P: Prophase, PM/M: Prometaphase/metaphase, A: Anaphase, T: Telophase.

See also Table S2.

Segú Cristina et al, Figure S6

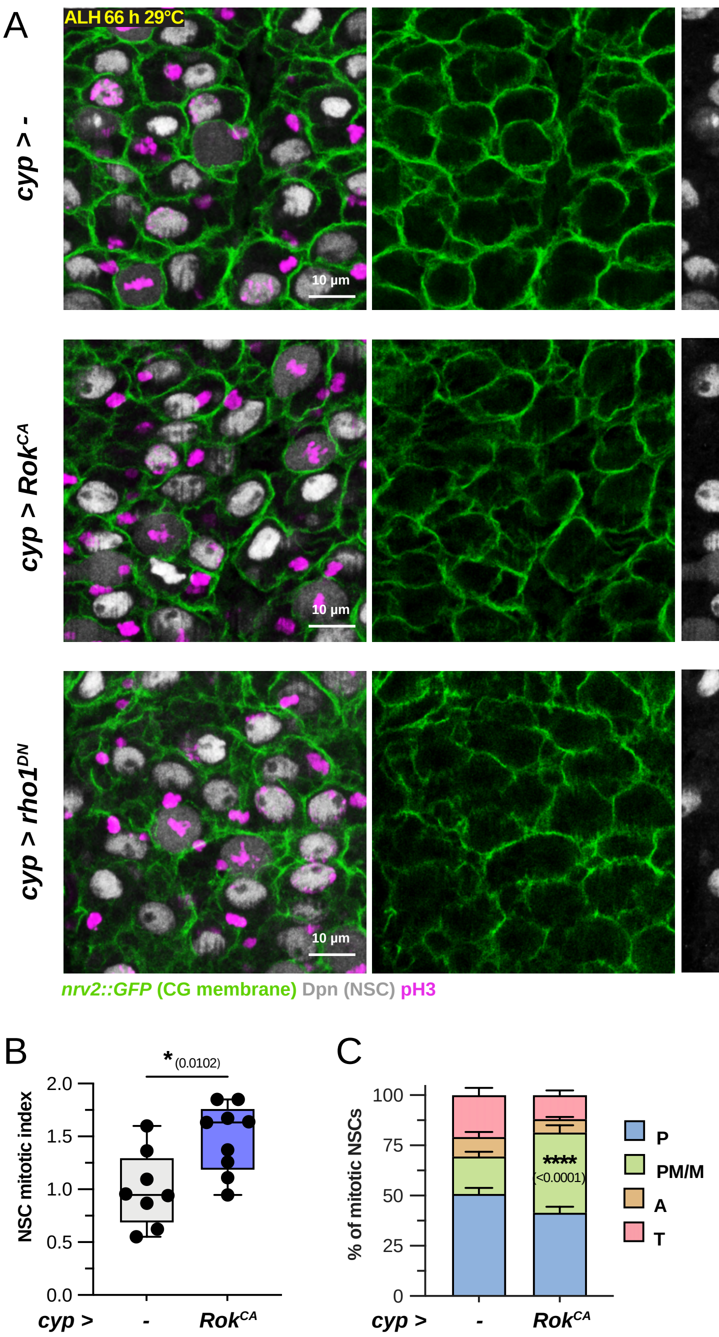

**Figure S6. Cortical tension in the cortex glia regulates neural stem cell nuclear shape and proliferation, related to Figure 5**

(A) Confocal images of the tVNC in control condition (*cyp* > -), constitutive activation of actomyosin contractility (*cyp* > $ROK^{CA}$) and constitutive inhibition of actomyosin contractility (*cyp* > $rho1^{DN}$). Activation and inhibition are performed for 24 h at 29°C at early L3 stage. Dashed yellow circles show examples of deformed nuclei.

(B-C) Box plot of mitotic index (normalized to the control condition) and stacked bar chart of phase distribution in NSCs for *cyp* > - (n = 8 tVNCs) and *cyp* > $ROK^{CA}$ (n = 9 tVNCs). Unpaired Student's t-test (B) and two-way ANOVA with Šídák multiple comparison test (C). This experimental replicate shows a higher mitotic index coupled with higher percentage of NSCs in prometaphase/metaphase.

ns, p≥0.05; *, p < 0.05; **, p <0.01; ***, p<0.001; ****, p<0.0001

P: Prophase, PM/M: Prometaphase/metaphase, A: Anaphase, T: Telophase.

See also Table S2.

Segú Cristina et al, Figure S7

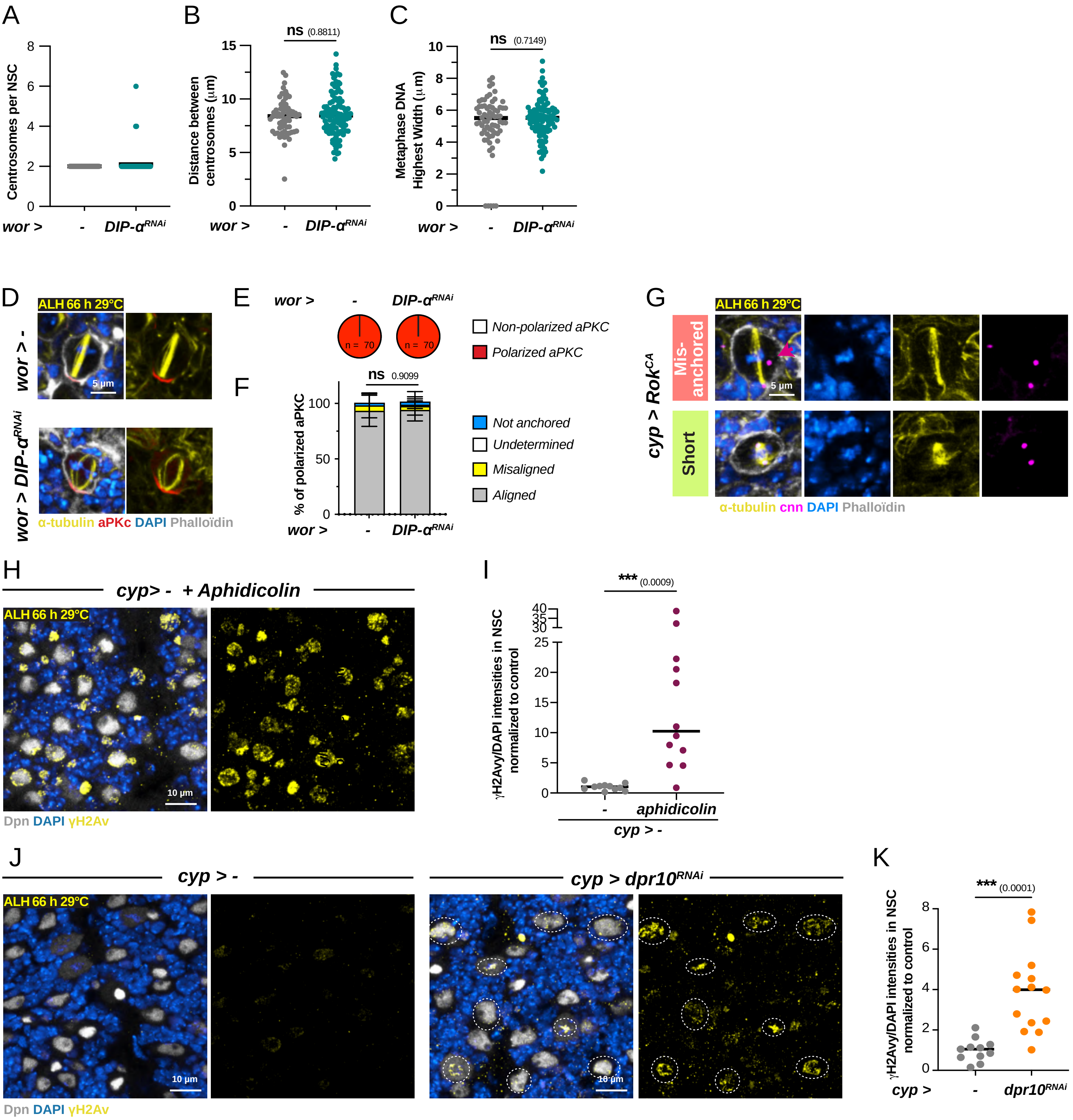

**Figure S7. IgSF-CAM$^{D3}$ disruption results in DNA damage and aneuploidy in neural stem cells, related to Figure 6**

(A) Dot plot of centrosome number per NSC in *wor* > - (n = 7 tVNCs and 75 NSCs) and *wor* > *DIP-α RNAi*$^{VDRC104044}$ (n = 9 tVNCs and 116 NSCs). Dot = NSC.

(B) Dot plot of the distance between centrosomes in NSCs in *wor* > - (n = 7 tVNCs and 67 NSCs) and *wor* > *DIP-α RNAi*$^{VDRC104044}$ (n = 9 tVNCs and 105 NSCs). Dot = NSC. Unpaired Student t-test.

(C) Dot plot of DNA width in metaphase NSC for *wor* > - (n = 7 tVNCs and 67 NSCs) and *wor* > *DIP-α RNAi*$^{VDRC104044}$ (n = 9 tVNCs and 105 NSCs). Dot = NSC. Mann-Whitney U test.

(D) Confocal images of NSC stained for the polarity marker aPKC (red) and the mitotic spindle (α-tubulin, yellow) in control (*wor* > -) and DIP-*α* knockdown (*wor* > *DIP-α RNAi*$^{VDRC104044}$) in NSCs. DAPI was used to counterstain the nuclei and Phalloidin the cellular boundaries.

(E) Pie charts of the distribution between polarized and non-polarized aPKC staining from D) for *wor* > - (n = 70 metaphase NSCs) and *wor* > *DIP-α RNAi*$^{VDRC104044}$ (n = 70 metaphase NSCs).

(F) Stacked bar chart of the percentage between aligned, misaligned, undetermined and non-anchored (spindle's pole not anchored to cortical aPKC) mitotic spindles (with respect to aPKC polarization) per tVNC for *wor* > - (n = 10 tVNCs) and *wor* > *DIP-α RNAi*$^{VDRC104044}$ (n = 10 tVNCs). Two-way ANOVA with a Dunnett's multiple comparison test. There is no significant difference for any of the three classes.

(G) Examples of abnormal mitotic spindles under constitutive activation of actomyosin contractility (*cyp* > *ROK*$^{CA}$). Pink arrowhead indicates a centrosome not aligned nor attached to the mitotic spindle.

(H) Confocal images of NSCs stained for the DNA damage marker γH2Av (yellow) in control (*cyp* > -) treated with Aphidicolin for 45 minutes.

(I) Dot plot of γH2Av intensity per NSC (normalized to DAPI intensity and then to control) in *cyp* > - (n = 11 tVNCs and 1718 NSCs) and *cyp*> - +Aphidicolin (n = 12 tVNCs and 1619 NSCs). Dot = tVNC. Unpaired Student t-test on the average values per tVNC.

(K) Confocal images of NSCs stained for the DNA damage marker γH2Av (yellow) in control (*cyp* > -) and *dpr10* knockdown in CG (*cyp*> *dpr10 RNAi*$^{VDRC103511}$).

(L) Dot plot of γH2Av intensity per NSC (normalized to DAPI intensity and then to control) in *cyp* > - (n = 11 tVNCs and 1718 NSCs) and *cyp*> *dpr10 RNAi*$^{VDRC103511}$ (n = 14 tVNCs and 2368 NSCs). Dot = tVNC. Unpaired Student t-test on the average values per tVNC.

For box plots, individual values are superimposed

ns, $p \geq 0.05$; *, $p < 0.05$; **, $p < 0.01$; ***, $p < 0.001$; ****, $p < 0.0001$

See also Table S2.

Segú Cristina et al, Figure S8

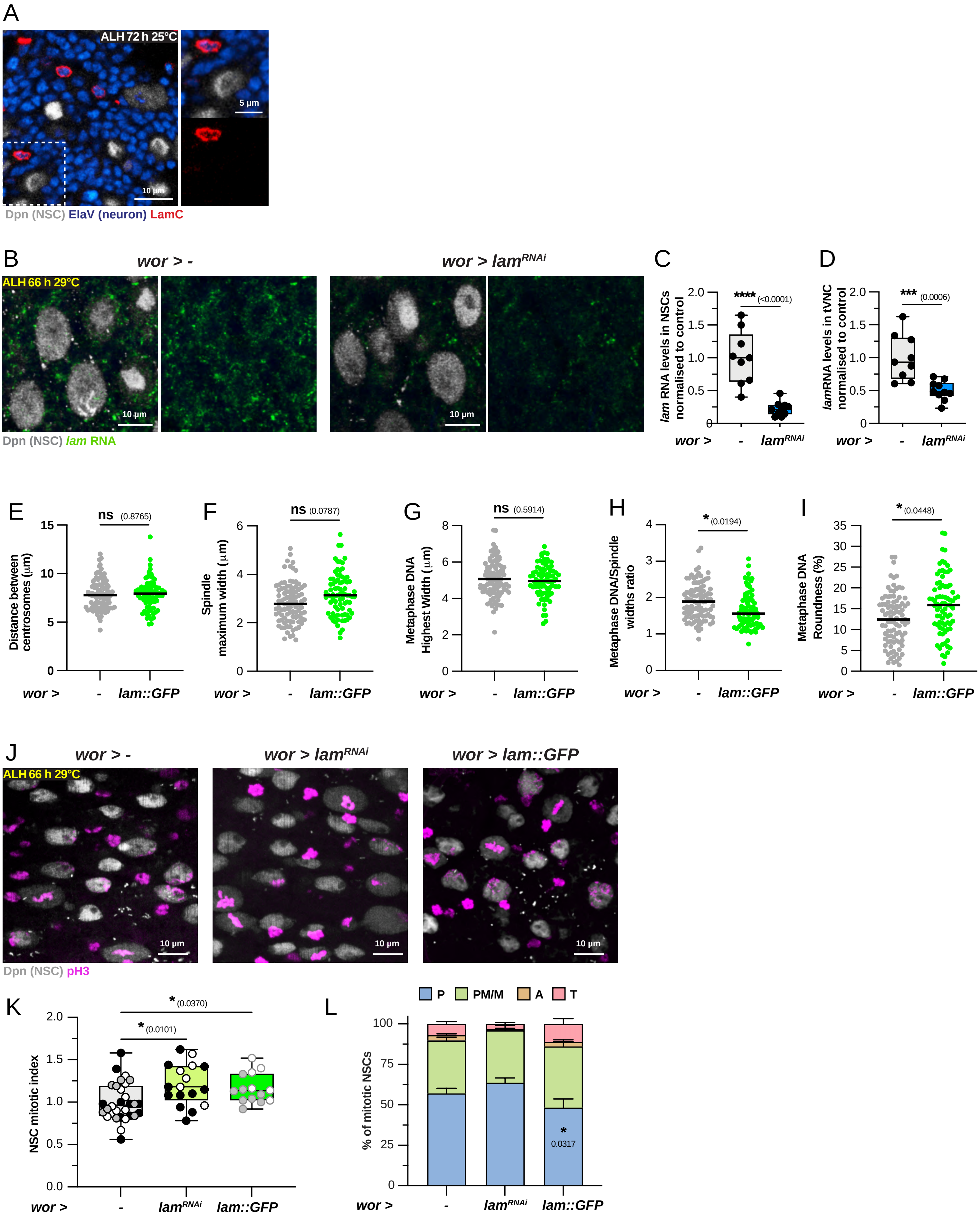

**Figure S8. Type-B lamin Lam modulates neural stem cell division, related to Figure 7**

(A) Confocal image of NSCs stained for the Type C lamin LamC (red) in wild-type tVNCs.

(B) Confocal images of a close-up of tVNCs stained for *Lam* mRNA for control (*wor*>-) and lamin knockdown (*wor*> $lam^{RNAi}$).

(C-D) Box plots of RNA levels in NSCs C) and tVNC D) (normalized to control) for *wor*>- (n = 9 tVNCs) and *wor*> $lam^{RNAi}$ (n = 10 tVNCs). Unpaired Student's t-tests.

(E) Dot plot of the distance between centrosomes in NSCs in *wor* > - (n = 11 tVNCs and 94 NSCs) and *wor* > *lam::GFP* (n = 12 tVNCs and 79 NSCs). Dot = NSC. Nested t-test on average values per tVNC.

(F-G) Dot plot of spindle (F) and DNA (G) widths for *wor* > - (n = 11 tVNCs and 94 NSCs) and *wor* > *lam::GFP* (n = 12 tVNCs and 79 NSCs). Dot = NSC. Nested t-tests on average values per tVNC.

(H) Dot plot of the ratio between DNA and spindle widths for *wor* > - (n = 11 tVNCs and 94 NSCs) and *wor* > *lam::GFP* (n = 12 tVNCs and 79 NSCs). Dot = NSC. Nested t-test on average values per tVNC.

(I) Dot plot of DNA roundness for *wor* > - (n = 11 tVNCs and 94 NSCs) and *wor* > *lam::GFP* (n = 12 tVNCs and 79 NSCs). Dot = NSC. Nested t-test on average values per tVNC.

(J) Confocal images of a close-up of tVNCs for control (*wor*>-), lam knockdown (*wor*> $lam^{RNAi}$) and lam overexpression (*wor*> *lam::GFP*) in NSCs showing NSC proliferation.

(K-L) Box plot of mitotic index (normalized to control) and stacked bar chart of phase distribution in NSCs for *wor* > - (n = 29 tVNCs), *wor*> $lam^{RNAi}$ (n = 17 tVNCs) and *wor* > *lam::GFP* (n = 14 tVNCs). Kruskal–Wallis H test with Dunn's multiple comparisons test (K) and two-way ANOVA with Šídák multiple comparison test (L).

For box plots, individual values are superimposed and colour-coded to represent multiple experimental replicates.

ns, $p \geq 0.05$; *, $p < 0.05$; **, $p < 0.01$; ***, $p < 0.001$; ****, $p < 0.0001$

P: Prophase, PM/M: Prometaphase/metaphase, A: Anaphase, T: Telophase.

See also Table S2.
