## Supplemental Tables for "Adhesion-Controlled Mechanics of the Glial Niche Regulate Neural Stem Cell Proliferative Potential": Table_S2.pdf

### Table S2. *Drosophila* genotypes, crosses and culture regimens per experiment

| Figures | Genotypes and crosses | Regimen |
|---|---|---|
| 1A | Nrv2::GFP/CyO; grh$^{D4}$-myr::mScarlet<br>x UAS-His2B ::YFP ; cyp4g15-GAL4 | ALH72 at 25°C |
| 1C-F | tub-GAL80$^{ts}$; cyp4g15-GAL4<br>x UAS-Dam<br>x UAS-Dam::PolII | T1: AEL32 at 18°C + 18h at 29°C<br>T2: 18°C until larval hatching (ALH0)<br>+ 32 h at 18°C + 18 h at 29°C<br>T3: 18°C until larval hatching (ALH0)<br>+ 76 h at 18°C + 18 h at 29°C |
| 2A-E<br>3H-K<br>4E-H<br>5A-C<br>S2B-F<br>S3A<br>S5B-E<br>S7H-K | Nrv2::GFP, tub-GAL80$^{ts}$ /CyO; cyp4g15-GAL4<br>x UAS-dpr10 RNAi$^{VDRC103511}$<br>x UAS-dpr6 RNAi$^{VDRC41161}$<br>x UAS-dpr6 RNAi$^{VDRC41161}$; UAS-dpr10 RNAi$^{BDSC27991}$<br>x w$^{1118}$ | 18°C until larval hatching (ALH0)<br>+ 66 h at 29°C |
| 2F-G | Nrv2::GFP, His-2Av::RFP/CyO; cyp4g15-GAL4<br>x UAS-dpr10 RNAi$^{VDRC103511}$<br>x w$^{1118}$ | 18°C until larval hatching (ALH0)<br>+ 66 h at 29°C |
| 3A-D | grh$^{D4}$-FLP; UAS-CD4::spGFP1-10, LexAop-CD4::spGPP11; cyp4g15-GAL4<br>x lexAop-myr::Tomato; grh$^{D4}$-FRT-stop-FRT-LexA<br>x dpr10 RNAi$^{VDRC103511}$, lexAop-myr::Tomato; grh$^{D4}$-FRT-stop-FRT-LexA | 18°C until larval hatching (ALH0)<br>+ 66 h at 29°C |
| 3L-M<br>6A-F, J-M<br>7A-C<br>S3C<br>S7A-F | worniu-GAL4, tub-GAL80$^{ts}$/CyO, GFP<br>x UAS-DIP-$\alpha$ RNAi$^{VDRC104044}$<br>x w$^{1118}$ | 18°C until larval hatching (ALH0)<br>+ 66 h at 29°C |
| 3N | mKate2::dpr10 x Nrv2::GFP/CyO | ALH72 at 25°C |
| 3O | DIP-$\alpha$::GFP x grh$^{D4}$-myr::mScarlet | ALH72 at 25°C |
| 3P | mKate2::dpr10 x DIP-$\alpha$::GFP | ALH72 at 25°C |
| 4A-D<br>S4A-D<br>S6A-G,L-M | Nrv2::GFP, worniu-GAL4/CyO; tub-GAL80$^{ts}$<br>x UAS-DIP-$\alpha$ RNAi$^{VDRC104044}$<br>x w$^{1118}$ | 18°C until larval hatching (ALH0)<br>+ 66 h at 29°C |
| 5D-G<br>7D-F<br>S6A-C | Nrv2::GFP, tub-GAL80$^{ts}$ /CyO; cyp4g15-GAL4<br>x UAS-ROK$^{CA}$<br>x UAS-rho1$^{DN}$<br>x w$^{1118}$ | 18°C until larval hatching (ALH0)<br>+ 72h at 18°C<br>+ 24 h at 29°C |
| 5H-K | NP577; Nrv2::GFP/CyO; dpr10$^{null}$/TM6,Tb<br>x UAS-rho1$^{DN}$; tub-GAL80$^{ts}$;dpr10$^{null}$/TM6,Tb<br>x tub-GAL80$^{ts}$;dpr10$^{null}$/TM6,Tb<br>x w$^{1118}$ | 18°C until larval hatching (ALH0)<br>+ 72h at 18°C<br>+ 24 h at 29°C |
| 6G-I<br>S7G | tub-GAL80$^{ts}$ /CyO; cyp4g15-GAL4<br>x UAS-ROK$^{CA}$<br>x w$^{1118}$ | 18°C until larval hatching (ALH0)<br>+ 72h at 18°C<br>+ 24 h at 29°C |

| | | |
|---|---|---|
| 7G-H, K-L<br>S8E-I | *worniu-GAL4, tub-GAL80$^{ts}$/CyO, GFP*<br>*x UAS-lamin::GFP*<br>*x $w^{1118}$* | 18°C until larval hatching (ALH0)<br>+ 72h at 18°C<br>+ 24 h at 29°C |
| 7I-J, M-O | *worniu-GAL4, tub-GAL80$^{ts}$/CyO, GFP*<br>*x UAS-DIP-α RNAi$^{VDRC104044}$; UAS-mCherry RNAi*<br>*x UAS-DIP-α RNAi$^{VDRC104044}$; UAS-lamin RNAi*<br>*x UAS-DIP-α RNAi$^{VDRC104044}$; UAS-lamin::GFP*<br>*x $w^{1118}$* | 18°C until larval hatching (ALH0)<br>+ 66 h at 29°C |
| S1B | *Nrv2::GFP/CyO* | T1: AEL32 at 18°C + 18h at 29°C<br>T2: 18°C until larval hatching (ALH0)<br>+ 32 h at 18°C + 18 h at 29°C<br>T3: 18°C until larval hatching (ALH0)<br>+ 76 h at 18°C + 18 h at 29°C |
| S2A | *Nrv2::GFP/CyO* | ALH72 at 25°C |
| S3B, D | *UAS-His2B::RFP; UAS-mCD8::GFP*<br>*x dpr10-GAL4*<br>*x dpr6-GAL4*<br>*x DIPA-α-GAL4* | ALH72 at 25°C |
| S3E | *DIPA-α$^{NULL2}$/ DIPA-α$^{NULL2}$*<br>*grh$^{D4}$-myr::mScarlet* | ALH72 at 25°C |
| S3F | *Dpr10::GFP x cyp4g15-myr-mtd::Tomato* | ALH72 at 25°C |
| S3G | *Dpr10::GFP* | ALH72 at 25°C |
| S3H | *mKate2::Dpr10 x DIP-α::GFP* | ALH72 at 25°C |
| S4E | *Nrv2::GFP, tub-GAL80$^{ts}$ /CyO; cyp4g15-GAL4*<br>*x UAS-DIP-α RNAi$^{VDRC104044}$*<br>*x $w^{1118}$* | 18°C until larval hatching (ALH0)<br>+ 66 h at 29°C |
| S4F-H | *ELAV$^{C155}$-GAL4; Nrv2::GFP, tub-GAL80$^{ts}$ /CyO*<br>*x UAS-DIP-α RNAi$^{VDRC104044}$*<br>*x $w^{1118}$* | 18°C until larval hatching (ALH0)<br>+ 66 h at 29°C |
| S4I-K | *DIPA-α$^{NULL2}$/ DIPA-α$^{NULL2}$*<br>*DIPA-α$^{NULL2}$/ DIPA-α$^{NULL2}$ x $w^{1118}$*<br>*DIPA-α$^{NULL2}$/ DIPA-α$^{NULL2}$ x DIPA-α ::GFP* | ALH72 at 25°C |
| S5A | *Nrv2::GFP, worniu-GAL4/CyO; tub-GAL80$^{ts}$*<br>*x UAS-dpr10 RNAi$^{VDRC103511}$*<br>*x UAS-dpr6 RNAi$^{VDRC41161}$*<br>*x $w^{1118}$* | 18°C until larval hatching (ALH0)<br>+ 66 h at 29°C |
| S5F-I | *Nrv2::GFP, tub-GAL80$^{ts}$ /CyO; cyp4g15-GAL4*<br>*x UAS-dpr10 RNAi$^{BDSC27991}$*<br>*x $w^{1118}$* | 18°C until larval hatching (ALH0)<br>+ 66 h at 29°C |
| S8A | *$w^{1118}$* | ALH72 at 25°C |
| S8B-D, J-L | *worniu-GAL4, tub-GAL80$^{ts}$/CyO, GFP*<br>*x UAS-lamin RNAi*<br>*x UAS-lamin::GFP*<br>*x $w^{1118}$* | 18°C until larval hatching (ALH0)<br>+ 66 h at 29°C |
